# Deep profiling of plant stress biomarkers following bacterial pathogen infection with protein corona based nano-omics

**DOI:** 10.1101/2024.12.12.627884

**Authors:** Roxana Coreas, Nikitha Sridhar, Teng-Jui Lin, Henry J. Squire, Elizabeth Voke, Markita P. Landry

## Abstract

Detection and remediation of stress in crops is vital to ensure agricultural productivity. Conventional forms of assessing stress in plants are limited by feasibility, delayed phenotypic responses, inadequate specificity, and lack of sensitivity during initial phases of stress. While mass spectrometry is remarkably precise and achieves high-resolution, complex samples, such as plant tissues, require time-consuming and biased depletion strategies to effectively identify low-abundant stress biomarkers. Here, we bypassed these reduction methods via a nano-omics approach, where gold nanoparticles were used to enrich time- and temperature-dependent stress-related proteins through biomolecular corona formation that were subsequently analyzed by ultra-high performance liquid chromatography tandem mass spectrometry (UHPLC-MS/MS). This nano-omic approach was more effective than a conventional proteomic analysis using UHPLC- MS/MS for resolving biotic-stress induced responses at early stages of pathogen infection in *Arabidopsis thaliana*, well before the development of visible phenotypic symptoms, as well as in distal tissues of pathogen infected plants at early timepoints. The enhanced sensitivity of this nano-omic approach enables the identification of stress-related proteins at early critical timepoints, providing a more nuanced understanding of plant-pathogen interactions that can be leveraged for the development of early intervention strategies for sustainable agriculture.

## Introduction

Bacterial pathogen infections, which impact agriculture by decreasing both crop productivity and quality^1,2^, are expected to increase in specific regions of the world due to the impacts of climate change.^3^ The ability to rapidly detect and control pathogens in crops is crucial for maintaining food security and preventing the transmission of zoonotic pathogens into the human population. Viable pathogens in crops can be assessed using conventional methods including visual examination and culture isolation, the latter of which is labor- and resource-intensive.^4^ Additionally, while visual examinations are limited by subjective analysis and lack of sensitivity, especially for pathogen detection in early stages, whereas culture-based methods are prone to contamination and are unsuitable for unculturable pathogens. Recently, molecular methods, such as PCR and ELISA, have been used to detect pathogens in plants however, these methods can produce false positives due to their reliance on immunological or genetic markers, and specificity is largely dependent on targeting a known pathogen and/or gene.^4,5^ The identification of stress induced small molecules, such as reactive oxygen species (ROS), typically relies on chromogenic and chemiluminescent substrates, staining and microscopy to evaluate cytotoxicity, electrolyte leakage to analyze cell membrane integrity, or light-emitting chlorophyll fluorometers to measure photosynthetic efficiencies.^6,7^ However, these approaches lack sensitivity, specificity, and rely on semi-quantitative methods that are typically inefficient for stress detection in plants prior to evident phenotypic responses. Recently, nanotechnology has been developed to assess early indicators of plant stress through nano-sensors that detect small molecules, such as H_2_O_2_,^8^ salicylic acid,^9^ and extracellular adenosine triphosphate^10^, in real-time following the induction of abiotic and biotic stressors. However, these nano-sensors are limited to detecting 1 small molecule or phytohormone at a time and their multiplexing is currently not feasible. Lastly, mass spectrometry (MS) can probe changes at the proteomic and metabolomic levels in plants exposed to stress conditions,^11,12^ however the detection of low abundance biomarkers, which are critical for assessing and monitoring stress in plants and crops,^13^ remains a significant challenge^14,15^.

Nanotechnology has also been used to deep profile proteomes via an approach termed ‘nano-omics’, where nanoparticles are introduced into biological milieus as a diagnostic tool for the detection of diseases in complex fluids and tissues.^16–19^ Nano-omics utilizes the physicochemical properties of nanoparticles, specifically high surface-to-volume ratios and facility for surface functionalization, to rapidly enrich low-abundance biomolecules from complex samples for downstream analysis with MS, alleviating bottlenecks associated with complicated and time-consuming sample preparation methods. This unique analytical approach is founded on the concept of the biomolecular corona;^20,21^ biomolecules spontaneously adsorb onto the surfaces of nanomaterials when they are introduced into complex environments. The constituents of the biomolecular corona are comprised of biomolecules that possess strong affinity to the nanoparticles, and often contain biomolecules that are not abundant in the native biofluid, thus permitting their selective detection. Protein corona constituents are subsequently characterized with analytical tools, such as MS. Moreover, relative to conventional MS-based liquid biopsies or sampling of biological milieus, such as plant leaf lysates, nano-omics reduces the need for extensive extraction, purification, and depletion strategies traditionally used to reduce the levels of high abundance proteins,^16^ thereby streamlining the detection of biomarkers. Importantly, stress biomarkers are inherently low-abundance biomolecules, especially in early timepoints following stress onset, thus making nano-omics a valuable approach for the early detection of stress-induced biomarkers in plants.

In this work, gold nanoparticles (AuNP), formulated with different surface charges, were used to enrich time- and temperature-dependent stress-related proteins from *Pseudomonas syringae* pathovar *tomato* (*Pst*) strain (DC3000) infected *Arabidopsis thaliana* Col-0 ecotype (hereafter referred to as *P. syringae* infected *A. thaliana*). AuNP were introduced into *A. thaliana* leaf lysates, and the resulting biomolecular corona was analyzed via nano-omics with ultra-high performance liquid chromatography tandem mass spectrometry (UPHLC-MS/MS). This nano-omic approach was compared to conventional proteomic analysis and was found to be more efficient at enriching and detecting stress-induced proteins in pathogen infected plant leaves, and non-infiltrated distal leaves from pathogen infected plants. Moreover, this nano-omic approach enhanced the detection of stress induced biomarkers at early timepoints prior to symptomatic expression and the onset of phenotypic responses in healthy appearing *A. thaliana*, enabling detection of early-onset plant stress, and potentiating its use for the future detection of new low-abundance plant stress biomarkers.

## Results

### Characterization of AuNP

Citrate capped (cit-AuNP) and branched polyethylenimine (BPEI) conjugated AuNP (BPEI-AuNP) were characterized for size, morphology, and surface properties using DLS, ζ-potential analysis, and UV-vis spectroscopy. DLS measurements conveyed that the hydrodynamic diameters of the cit-AuNP and BPEI-AuNP were 18.4 ± 1.4 and 39.6 ± 3.1 nm, respectively (**Supplementary Fig. 1A**). Correspondingly, ζ-potential measurements confirmed cit-AuNP had a negative charge of −40.6 ± 2.5 mV and BPEI-AuNP had a positive charge of 36.3 ± 1.7 mV at pH 6 in Milli-Q water (18.2 MΩ/cm) (**Supplementary Fig. 1B**), supporting the presence of the negatively charged citrate ligand and the positively charged BPEI polymer on the AuNP surfaces. The UV-vis spectra of cit-AuNP exhibited an adsorption band with a λ_max_ of 519.5 nm, which is characteristic of spherical AuNPs that are stable in solution.^22^ Conjugation with BPEI resulted in a slight red shift (BPEI-AuNP λ_max_ 524.5 nm) (**Supplementary Fig. 1C**). Average polydispersity index (PDI) values were 0.27 ± 0.01 and 0.16 ± 0.02 for BPEI-AuNP and cit-AuNP, respectively (**Supplementary Fig. 1D**), suggesting that the AuNP were moderately polydispersed.

### Deep profiling of time-dependent pathogen infected *A. thaliana* proteome with protein corona based nano-omics

To determine if pathogen induced stress markers could be detected prior to the manifestation of phenotypic expression of disease in *A. thaliana*, we applied a nano-omic strategy and compared its efficiency to a conventional proteomics approach. We used three plants as biological replicates for each condition and timepoint. For each plant, primary inoculation of *P. syringae* was performed by pressure infiltration into the apoplast of three 5-6 week old leaves distributed ∼130° apart.^23^ Mock treated plants were infiltrated with 10 mM MgCl_2_. A schematic representation and photographs of the temporal phenotypic effects observed in pathogen infected *A. thaliana* are shown in **Fig. 1A** and **Supplementary Fig. 2**, respectively. While 10 mM MgCl_2_ did not induce observable phenotypic changes in *A. thaliana* over time, exposure to *P. syringae* resulted in a time-dependent manifestation of chlorosis, leaf wilting, necrotic lesions, and stunted growth, aligning with previous studies,^24–26^ with severe symptoms occurring between 3- and 7-DPI (days post infiltration). At designated timepoints (0.5-, 1-, 3- and 7-DPI), six leaves were collected from each plant: three infiltrated leaves and three non-infiltrated ‘distal’ leaves. Leaves collected from 0.5- and 1-DPI plants were designated ‘early timepoint’ samples, while those collected from 3- and 7-DPI were assigned ‘late timepoint’. An additional control group of non-infiltrated plants was also sampled to determine how the mock treatment would impact the *Arabidopsis* proteome. For each condition and timepoint, we used three plants as biological replicates. The three leaves collected from each plant for each condition (infiltrated, distal, and non-infiltrated control) and timepoints were pooled and treated as a single biological sample. The collected leaves were briefly washed to sterilize the surface, flash frozen, and lysed in Milli-Q water using a bead-beater homogenizer. We then compared two approaches to analyze the proteome of *A. thaliana*: (1) a conventional proteomic analysis where the homogenized leaf lysates were directly analyzed by UHPLC-MS/MS, and (2) our nano-omic approach where biomolecular coronas were formed by incubating BPEI-AuNP and cit-AuNP with leaf lysates at ambient temperatures and the proteins from the coronas were analyzed with UHPLC-MS/MS. Both approaches followed a 10 minute electrophoretic separation step to resolve the proteins; this brief separation was applied to ensure separation quality with rapid processing of the samples.

**Fig. 1.**
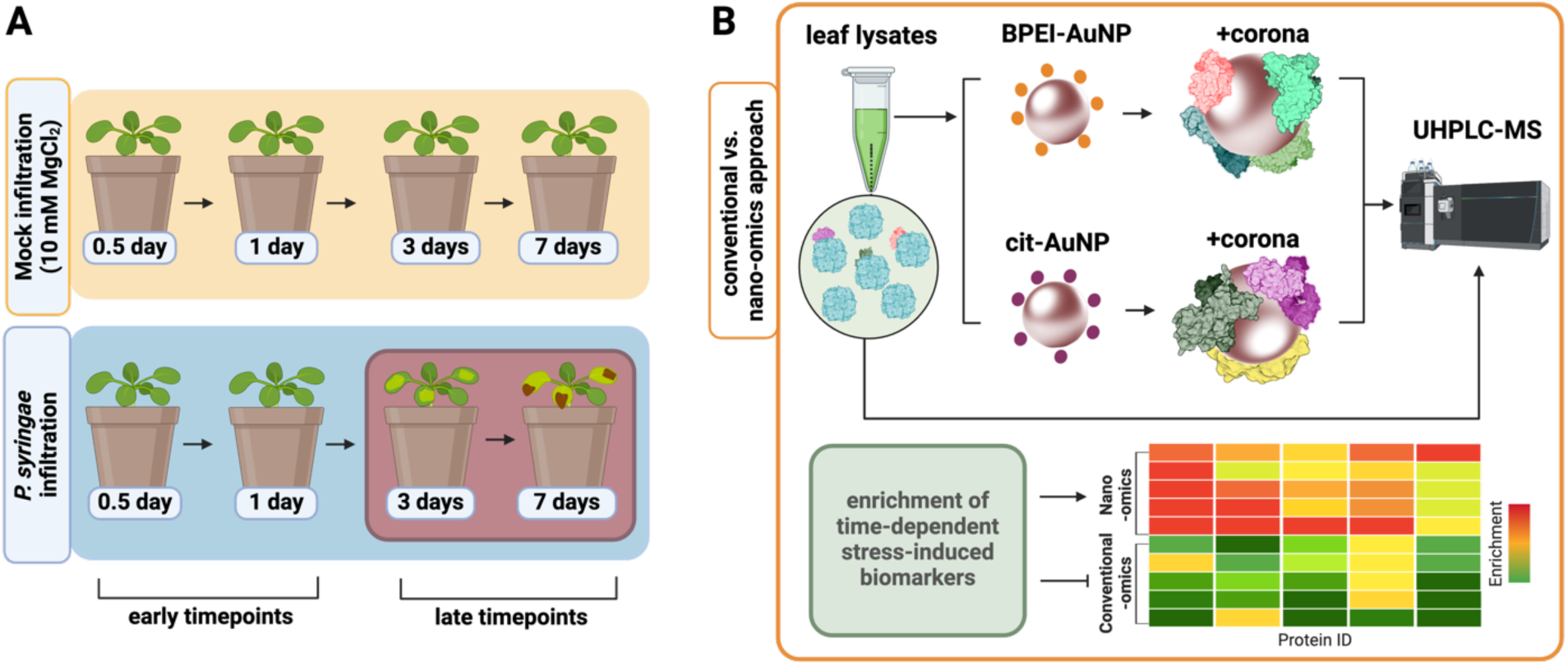
**A)** Scheme of the biotic stress induced in *A. thaliana* and the deep profiling nano-omics approach used in this study. Following pathogen infection, *A. thaliana* tissues, both directly infected leaves as well as distal leaves from infected plants, were collected from 0.5- to 7-DPI timepoints (blue box) for analysis. Mock treated plants were infiltrated with 10 mM MgCl_2_ (yellow box) and leaves were collected from 7-DPI mock treated plants as controls. Time-dependent phenotypic responses in pathogen infected *A. thaliana* were apparent at 3- and 7-DPI (red box). **B)** Protein markers of stress were probed in pathogen infected *A. thaliana* leaf lysates using conventional proteomic and nano-omic approaches. While both approaches employed UHPLC-MS/MS, nano-omics leveraged protein corona enrichment with BPEI-AuNP and cit-AuNP.

Between the conventional and nano-omic approaches, 3128 and 559 *A. thaliana* and *P. syringae* proteins were comprehensively identified. Average number of proteins and CV percentages per sample type are listed in **Supplementary Table 1**. Protein spectral counts were analyzed by principal component analysis (PCA) to reduce high-dimensional data into two principal components, with the resulting scores plots shown in **Fig. 2**. We observed overlaps and proximity clustering among samples that were analyzed by conventional proteomics. Specifically, the mock, 0.5-DPI, and 0.5-DPI distal samples overlapped, and were positioned near the non-infiltrated control samples (**Fig. 2A**). A similar proximity grouping was observed between the control, mock, 1-DPI and 1-DPI distal samples (**Fig. 2B**). In contrast, while the mock and 3-DPI distal samples also overlapped and grouped near the control samples, the 3-DPI group was positioned further away (**Fig. 2C**), indicating some divergence. Likewise, the control, mock, and 7-DPI distal samples were closely grouped together, but the 7-DPI samples were further removed, suggesting a greater degree of variance in the data collected at this timepoint (**Fig.2D**). These findings suggest that samples from the early and non-symptomatic timepoints (0.5- and 1-DPI) exhibit protein compositions that are highly similar to the mock infiltrated and non-infiltrated control plants, with minimal differences detectable by the conventional proteomic approach. Protein coronas extracted from the BPEI- and cit-AuNPs were positioned furthest away from the conventionally analyzed samples, with clear discrepancies between them. Coronas formed with the lysates from early timepoints (0.5- and 1-DPI) grouped closer together based on surface charge and not sample type (distal vs infiltrated) (**Figs. 2A, 2B**). In contrast, coronas formed with lysates from the late timepoints (3- and 7-DPI) tended to cluster together according to sample type (**Figs. 2C and 2D**), suggesting that late timepoints exhibit a larger degree of stress-induced changes in the proteome, therefore relatively diminishing the importance of AuNP surface chemistry for nano-omic analysis. Notably, overlaps were observed between coronas formed with cit-AuNPs and the early timepoint samples, suggesting that BPEI-AuNPs may be more effective than cit-AuNP in adsorbing more unique proteins from early-stage pre-symptomatic infected plants. Additionally, the cit-AuNP coronas formed with 3- and 7-DPI samples overlapped with the conventionally analyzed 3- and 7-DPI samples, indicating that the protein corona compositions formed with cit-AuNP closely resembled protein compositions in the native biofluid. In contrast, the BPEI-AuNP coronas displayed greater divergence from the native biofluid protein compositions. PCA was also conducted on coronas formed with the mock, mock distal, and non-infiltrated control samples as a subset analysis. As shown in **Supplementary Fig. 3**, no clear distinction between clusters was observed between the control, mock, mock distal lysates, as expected, and their corresponding cit-AuNP coronas, indicating a close similarity between these groups. BPEI-AuNP coronas formed with non-infiltrated control, mock control, and mock distal samples were positioned furthest away. Collectively, these results suggest that cit-AuNP may reflect the native biofluid protein composition more closely, while BPEI-AuNP could be more selective for distinct or ‘unique’ proteins. This selectivity, particularly in the context of disease progression, implies that BPEI- AuNP may capture unique proteomic signatures, or biomarkers, that are not as easily detected through conventional proteomics, and may do so more effectively than cit-AuNP. **Supplementary Fig. 4** presents the PCA plot encompassing all analyzed samples together. To quantify the divergence in PCA space between samples, the Euclidean distances of the mean PC1 and PC2 values were calculated for the time dependent samples relative to the mock, as shown in **Supplementary Fig. 5**. With increasing timepoints (0.5- to 7-DPI), we observed a progressive increase in the Euclidean distance from the mock for samples analyzed by conventional proteomics, with distances ranging from 2.9 – 24. A similar trend was observed for distal tissues analyzed by conventional proteomics, where distances ranged from 2 –17.6. Collectively, samples analyzed by the conventional proteomic approach were closest to the mock in PCA space, supporting that their protein profiles were most similar to the mock samples. In contrast, samples analyzed using nano-omics exhibited significantly greater Euclidean distances from the mock, with values ranging between 28 – 72. Among the AuNP coronas, those formed with the BPEI- AuNP showed the largest distinction from the mock, as evidenced by their greater Euclidean distances. Notably, with the nano-omics samples, we did not see a progressive increase in Euclidean distance over time. These findings highlight the sensitivity of the nano-omics approach in capturing protein differences across samples mediated by distinct differences in AuNP surface chemistry, and suggest that AuNP-based enrichment of low-abundance stress markers, especially at early disease timepoints, is a valuable method for detecting pre-symptomatic plant stress.

**Fig. 2.**
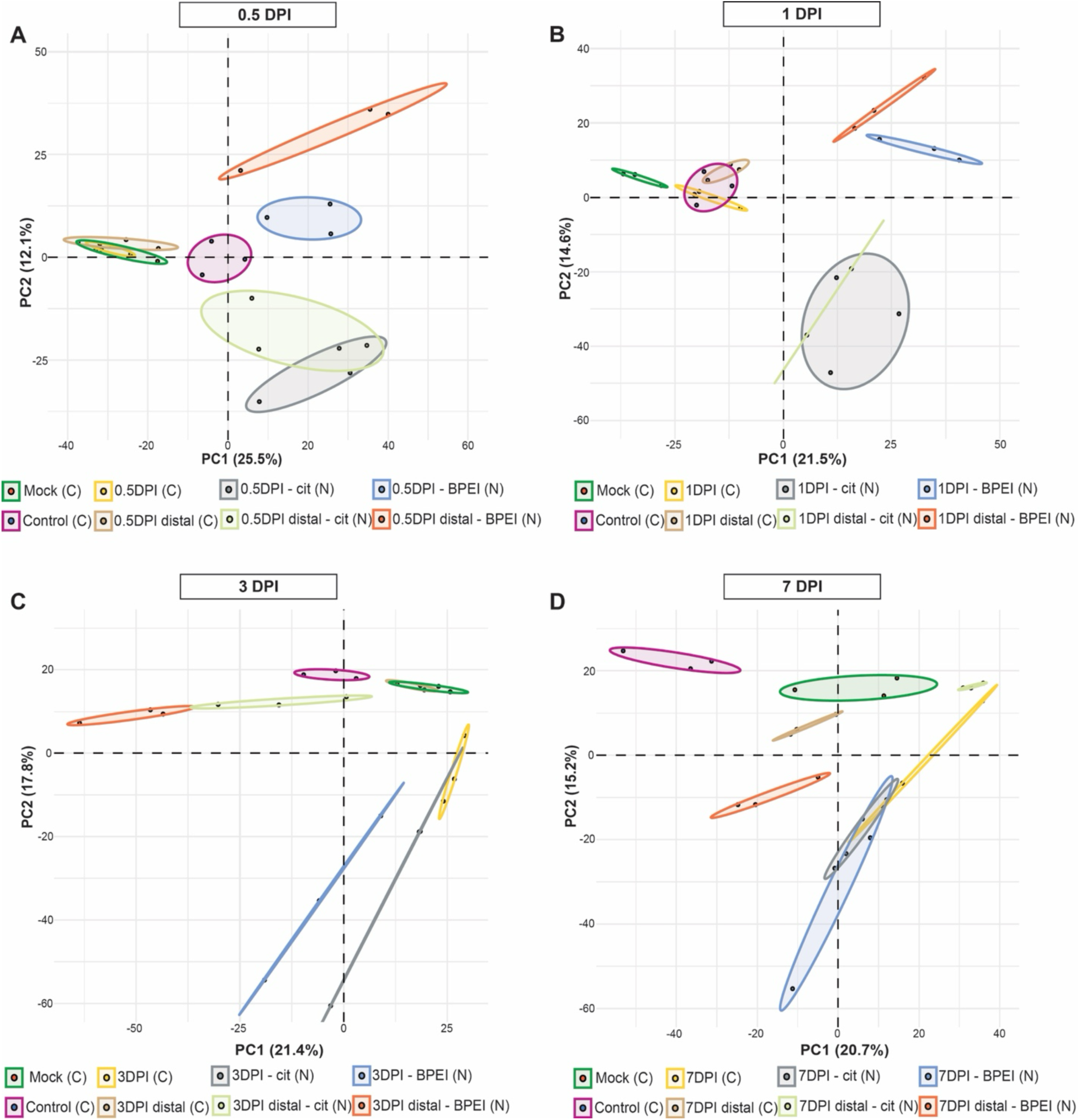
Principal component analysis of protein spectral counts identified by the conventional proteomic and nano-omic approaches. Score plots of the lysate, distal leaf lysate, BPEI-AuNP corona, cit-AuNP corona, mock, and non-infiltrated control for **A)** 0.5-DPI, **B)** 1-DPI, **C)** 3-DPI, and **D)** 7-DPI conditions; PC1 and PC2 summarized 37.6%, 36.1%, 39.2%, and 35.9% of the variances, respectively. Each point represents a biological replicate, and the ellipses represent the 95% confidence intervals around the mean point of each group (*n*=3). In **B)**, only 2 of the 3 biological replicates are shown and analyzed by PCA for the 1-DPI distal-cit-AuNP group; the PCA of all three replicates for the 1-DPI distal-cit-AuNP group is shown in **Supplementary Fig. 6.** (C) denotes that the sample was analyzed by conventional proteomics and (N) denotes that the sample was analyzed by the nano-omic approach.

Z-scores, calculated from the average relative abundance of proteins, are displayed as heatmaps with hierarchical clustering in **Supplementary Figs. 7, 8** for *A. thaliana* and *P. syringae* proteins, respectively. For *A. thaliana* proteins, clustering of z-scores revealed patterns of similarity between lysates and coronas that were dependent on timepoint, sample type (infiltrated, mock, distal, non-infiltrated control) and AuNP surface chemistry. Z-score analysis suggests that while the surface chemistry of the AuNP plays a dominant role in protein corona formation at earlier timepoints, its influence diminishes as disease progresses. Importantly, this analysis shows greater variability in protein samples derived from AuNP-based corona samples relative to those analyzed by conventional proteomics, suggesting that the nano-omic approach captured a wider range of *A.thaliana* proteins with relative changes in expression levels. Among the samples from the later timepoints (3- and 7-DPI), which exhibited the greatest variation in protein expression, the AuNP coronas had higher variability compared to the corresponding lysates analyzed by conventional proteomics, suggesting that the nano-omic approach can provide more nuanced insights on pathogen infection.

To further compare relative protein expression, log_2_(fold changes) were calculated by comparing protein levels in pathogen-infiltrated leaf lysates and the AuNP coronas formed with these lysates against those in mock treated and non-infiltrated controls. Additionally, proteins exclusively detected in either the lysates or the AuNP coronas – but absent from the mock treated and/or non-infiltrated controls – were classified as ‘unique’ proteins. **Supplementary Fig. 9** illustrates the quantity of differentially expressed (log_2_(fc) >1 or <-1) and ‘unique’ *A. thaliana* and *P.syringae* proteins identified, relative to protein expression in mock treated leaf lysates, via the nano-omic and conventional proteomic approaches. Across the timepoints and sample types (distal and infiltrated), the nano-omics approach allowed for the characterization of 1.3 – 3.7 fold more proteins compared to conventional proteomics. To illustrate the relationship between these proteins and the analytical approaches applied, Venn diagrams were generated to compare protein profiles across three time-dependent conditions: cit-AuNP corona proteins, BPEI-AuNP corona proteins, and proteins in the leaf lysates analyzed through conventional proteomics. In general, the number of *A. thaliana* proteins exclusive to each type of AuNP was larger than the quantity of proteins identified by conventional proteomic analyses. However, this trend was less prominent for *P. syringae* proteins, particularly in early timepoint infiltrated samples and distal leaf lysates, where fewer proteins were observed. Neither of the two AuNP surface chemistries used in this study appeared to be more efficient than the other at enriching *P.syringae* proteins. Similarly, **Supplementary Fig. 10** depicts the number of differentially expressed (log_2_(fc) >1 or <-1) and ‘unique’ *A. thaliana* and *P.syringae* proteins identified relative to non-infiltrated controls, where the nano-omics approach facilitated the identification of 1.3 – 3.8 fold more proteins than the conventional proteomic analyses. Furthermore, Venn diagrams revealed that protein enrichment, specifically the number of proteins, on the cit-AuNP and BPEI-AuNP was not strongly influenced by surface chemistry, suggesting that both nanomaterial types can capture a broad range of proteins regardless of surface properties.

The shared and distinct proteins identified between differentially expressed and ‘unique’ proteins were then used to calculate Jaccard index values to quantify the degree of similarity across samples. These similarity values were plotted as heatmaps with hierarchical clustering, as shown in **Fig. 3**. For pathogen infiltrated leaves, the similarity heatmap of *A. thaliana* proteins revealed two distinct clusters that separate the conventional proteomic analyses from the nano-omics approach. Sub-clustering of the nano-omics samples revealed distinct grouping patterns: for the late timepoints (3- and 7-DPI), samples clustered according to sample type, while for early timepoints (0.5- and 1-DPI), subculturing was driven by the AuNP surface chemistry. Meanwhile, the similarity heatmap for *P. syringae* proteins exhibited grouping that was driven by time-dependence, with samples from early timepoints (0.5- and 1-DPI) clustering together and separately from later timepoints (3- and 7-DPI). Congruently, Jaccard index values were also calculated for *A. thaliana* and *P. syringae* proteins identified from distal leaves of pathogen infected plants (**Fig. 3**). Notably, the degree of similarity of *A.thaliana* proteins across distal samples was generally higher than their similarity in pathogen infiltrated leaves. In contrast, *P. syringae* proteins exhibited a more complex pattern of similarity. An opposing trend was observed, where the similarity between distal leaf samples was comparatively lower than in the pathogen-infiltrated samples. To compare protein similarities with mock infiltrated plants, we calculated Jaccard index values by comparing the differential expression and ‘unique’ protein composition of each sample relative to non-infiltrated controls (**Supplementary Fig. 11**). This approach enabled a quantitative evaluation of how protein compositions in our samples diverged from those in healthy, non-infiltrated control plants. Hierarchical clustering revealed that mock treated samples were most similar to the 0.5-DPI samples, regardless of sample type (distal vs infiltrated) or protein origin (*A. thaliana* vs *P.syringae*). Additional clusters reflected the same trends driven by time-dependence and AuNP surface chemistry, consistent with the patterns observed in **Fig. 3**. This clustering pattern underscores that early-stage protein expression in pathogen infiltrated plants closely resembles that of mock treated plants and this similarity may reflect the pathogen’s initial phase of growth, during which it has not yet reached a sufficient population to elicit a pronounced differential response in the plant, thus a conventional proteomic approach may not be sensitive enough to distinguish low-abundant stress-induced proteins at early and critically relevant timepoints.

**Fig. 3.**
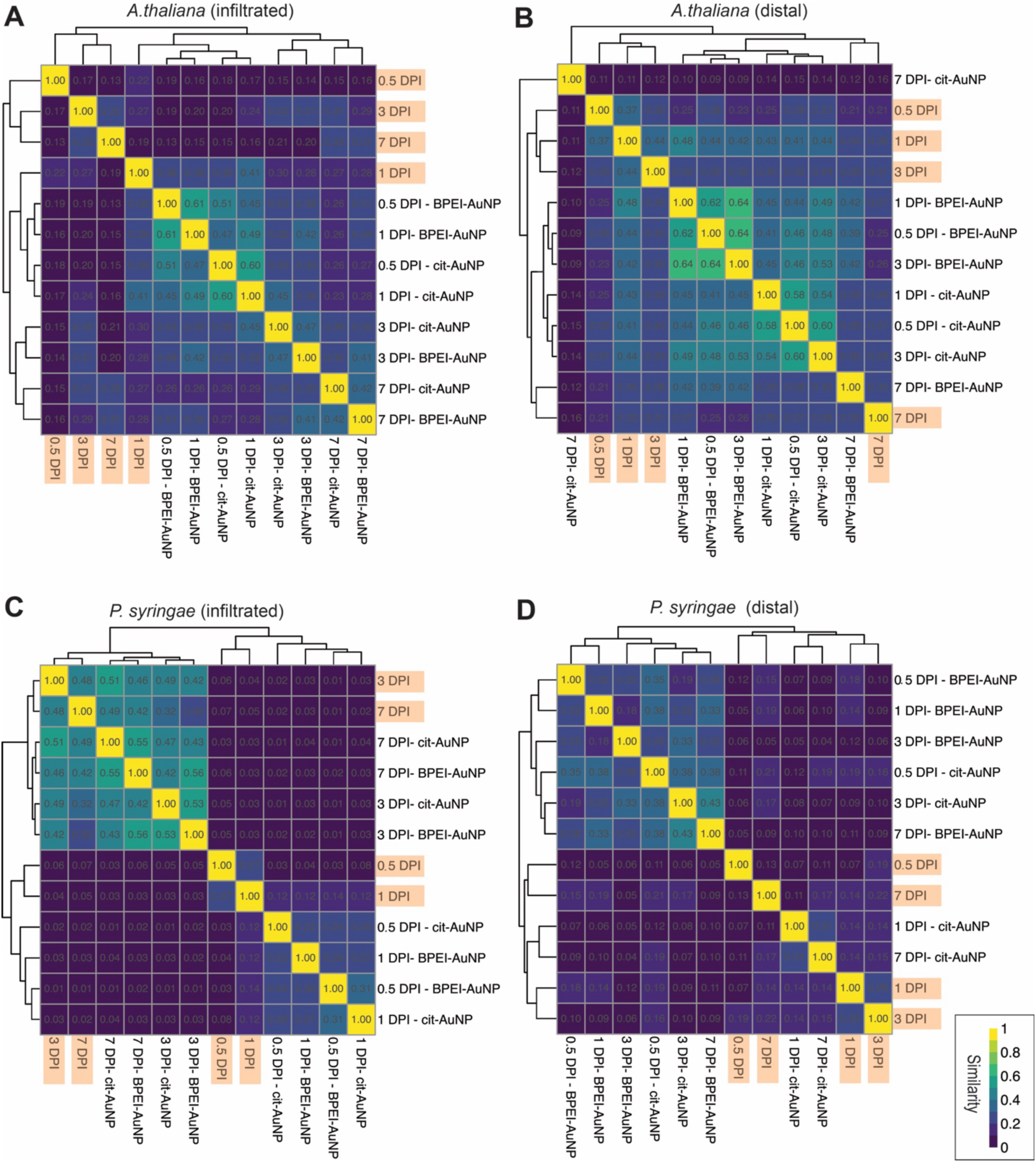
Heatmaps with hierarchical clustering depict the similarities between samples based on the composition of **A), B)** *A. thaliana* and **C), D)** *P. syringae* differentially expressed (log_2_(fc)>1 and <-1) and unique proteins, relative to mock infiltration, identified in **A), C)** pathogen infiltrated and **B), D)** distal tissues of pathogen infected plants, respectively. The color scale indicates the degree of similarity based on Jaccard index values. Samples labeled in orange represent those analyzed by the conventional proteomic approach.

To better highlight protein patterns in response to pathogen infiltration, we generated heatmaps of differential expression (i.e.: log_2_(fc) values), comparing protein levels in lysates and AuNP coronas against those in mock infiltrated samples. This approach prioritizes variations in protein abundances, revealing over- and under-expressed proteins in lysates analyzed via the conventional proteomic approach, as well as enriched and depleted proteins in the AuNP coronas analyzed via nano-omics. **Fig. 4A** shows the heatmap of differential protein expression in pathogen-infiltrated lysates, while **Fig. 4B** shows protein expression in lysates of distal leaves from pathogen-infiltrated plants. For both heatmaps, proteins were filtered to include only those that met the conditions of log_2_(fc) >3.5 and *p*<0.05 in at least one sample. We identified four proteins associated with bacterial infection responses that were significantly enriched on the AuNPs. Nodulin/glutamine synthetase-like protein (NodGS) (F4J9A0), which can also be involved in developmental processes independent of bacterial infections,^27^ was slightly overrexpressed following pathogen infection according to conventional proteomics, but was significantly enriched on the BPEI-AuNP corona formed with the 0.5-, 1-, and 3-DPI lysates, suggesting that this stress biomarker is better detected with nano-omics. Interestingly, in distal tissues, NodGS was significantly enriched on the BPEI-AuNP coronas formed with the 0.5-DPI lysates, but not detectable by conventional proteomics, indicating that, although distal tissues appeared healthy, their proteome suggests an underlying response to pathogen-induced signals from infected leaves. These findings imply that even tissues not directly exposed to the pathogen may initiate subtle defense mechanisms in response to systemic infection signaling. Ferritin-1 (Q39101), which regulates iron homeostasis and leaf development,^28^ slightly increased with pathogen infection, particularly in the late timepoint samples (3- and 7-DPI), as detected by conventional proteomics. Ferritin-1 was enriched on cit-AuNP, specifically in the coronas formed with 0.5- and 1-DPI, supporting its expression as an early, low abundance protein, induced by pathogen infection^29^ that is detectable at early timepoints solely by nano-omics. Lipoxygenase 2 (P38418), a jasmonate-inducible enzyme integral to various plant defense pathways,^30,31^ was notably overexpressed in pathogen-infiltrated samples, with peak expression observed at 7-DPI, supporting previous findings.^32^ Significant enrichment of this enzyme was also observed on both AuNP coronas, particularly at early infection stages (0.5- and 1-DPI). These findings indicate that lipoxygenase 2 may play an early and sustained role in the plant’s defense response, and AuNP coronas effectively capture and enrich this protein during initial pathogen exposure. Cinnamyl alcohol dehydrogenase 7 (Q02971), encoded by the *ELI3* gene and inferred to be overexpressed as result of pathogen infection,^33^ was only significantly overexpressed in 1-DPI lysates. It was significantly enriched on both AuNP coronas formed with the earliest timepoint samples (0.5-DPI), indicating that it may be more efficiently captured on the AuNP coronas than with conventional proteomics due to its low abundance in the 0.5-DPI lysates. Additionally, we observed the enrichment of several proteins associated with redox homeostasis in both pathogen infiltrated samples and lysates of distal leaves. Peroxisomal malate dehydrogenase 1 (O82399), along with acyl-coenzyme A oxidases 3 (P0CZ23) and 4 (Q96329), were more significantly enriched on AuNPs when compared to their expression levels in lysates analyzed by conventional proteomics. These enzymes play a key role in regulating fatty acid β-oxidation,^34,35^ a metabolic process that supports cellular redox homeostasis by generating electron carriers essential for maintaining oxidative stability under stress.^36^ Small ribosomal subunit proteins RACK1y (Q9C4Z6) and RACK1x (Q9LV28), MAPK cascade scaffolding proteins that regulate immune signaling pathways,^37^ were overexpressed and significantly enriched in 0.5-, 1-, and 3-DPI distal lysates and coronas. However, the detection of these proteins was more prominent through the nano-omic approach, especially for RACK1x which was resolvable in 7-DPI samples exclusively by the nano-omic approach. Moreover, several mRNA binding proteins were not significantly overexpressed in distal lysates when analyzed by conventional proteomics but showed selective enrichment on the BPEI-AuNP protein corona. For example, small ribosomal subunit protein eS6z (O48549) was significantly enriched on BPE- AuNP coronas formed with the 0.5-, 1-, 3- and 7-DPI distal lysates. While the expression of this protein in response to bacterial infections has not yet been characterized in *A. thaliana*, transcriptional upregulation of its gene has been documented in *Oryza sativa* (rice) in response to infection by the bacterial pathogen *Xanthomonas oryzae* pv. *oryzae*.^38^ This selective enrichment of eS6z and other mRNA binding proteins on BPEI-AuNP highlights the potential for nano-omic approaches to capture low-abundance, stress-responsive proteins that may be overlooked by conventional proteomics. Interestingly, *P.syringae* catalase-peroxidase (Q87WL6) was markedly overexpressed in 3- and 7-DPI lysates and their respective coronas. This finding is interesting as bacterial recovery from 7-DPI lysates was not feasible (**Supplementary Fig. 19**), suggesting that catalase-peroxidase expression may reflect a stress adaptation mechanism by *P.syringae* in response to the plant’s defenses. The sustained presence of catalase-peroxidase underscores its potential role in pathogen survival strategies within host tissues at advanced infection stages.

**Fig. 4.**
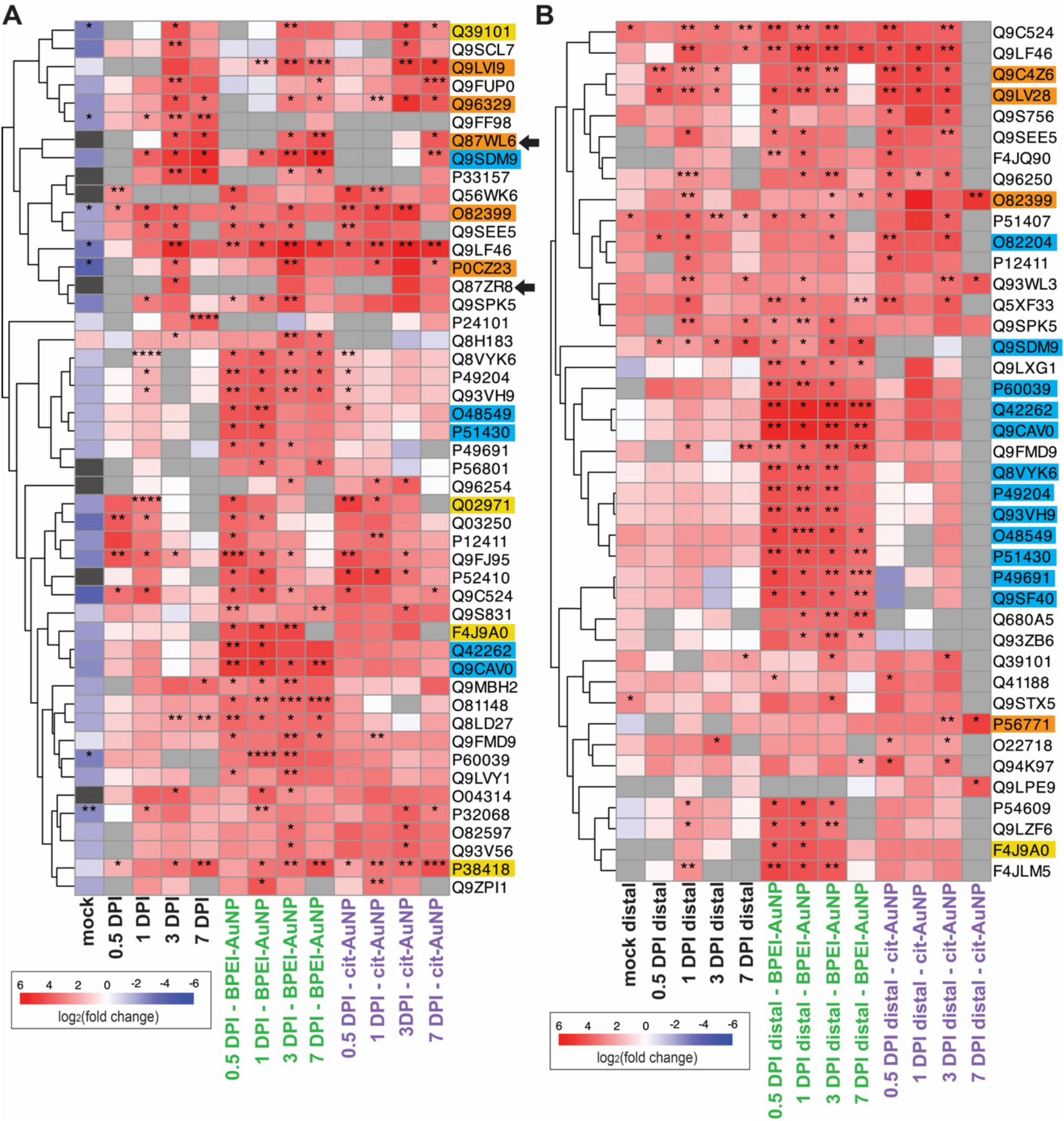
Heatmaps with Euclidean hierarchical clustering of a subset of log_2_(fc) values, calculated relative to mock infiltration, of proteins from **A)** pathogen infiltrated leaf lysates and **B)** distal tissues of pathogen infected plants. For the heatmaps, proteins were filtered to plot log_2_(fc) >3.5 and *p*-values<0.05 observed in at least one sample. Samples analyzed by conventional proteomics are labeled in black font, while nano-omics analyses are labeled in green and purple fonts for the BPEI- and cit-AuNPs, respectively. In panel **A),** mock refers to its protein expression relative to the non-infiltrated control. Overexpression, or protein enrichment on the AuNPs, is depicted by red boxes (log_2_(fc) >0), while underexpression, or depletion from the AuNP surfaces, is depicted by blue boxes (log_2_(fc) <0). Grey boxes indicate incalculable fold change. Light grey represents cases where the protein was detected solely in the mock, resulting in an undefined fc calculation. Dark grey represents cases where the protein was not detected in the mock, making the fc calculation invalid. Asterisks denote significance levels (*, *p*< 0.05; **, *p* <0.01; ***, *p*<0.001; ****, *p*<0.0001). Protein accession IDs are highlighted to annotate specific functions: proteins responsive to bacterial infection are marked with a yellow highlight; proteins involved with redox homeostasis are marked with an orange highlight; and proteins that bind to RNA are marked with a blue highlight. Black arrows point to *P.syringae* proteins.

To further determine if pathogen induced biomarkers were detectable in our samples, we identified enriched biological processes of the ‘unique’ plant proteins detected in plant tissues directly infected with *P.syringae*. The ‘unique’ proteins in our samples were those that were uniquely expressed in pathogen-infected tissues but were not identified in the mock treated controls. To analyze the enriched processes, we leveraged DAVID, a robust bioinformatics platform designed for gene ontology and pathway enrichment analysis.^39,40^ The unique proteins from the pathogen infected plants were associated with 260 enriched biological processes. Of these, 45% were identified solely by nano-omics, 17% were identified with conventional proteomics, and 38% were identified by both analytical approaches. We scrutinized the time-dependency of these enriched biological processes by categorizing them according to the earliest timepoint at which the highest gene count was observed. For the early timepoint samples (0.5- and 1-DPI), 43% (111) of the enriched biological processes were identified uniquely by nano-omics, 13% (33) by conventional proteomics, and 21% (56) were detected by both approaches; however, for those that were identified by both approaches, 19% (50) showed higher gene or proteins counts when analyzed by nano-omics. For the late timepoint samples (3- and 7-DPI), 11% (28) were detected solely by nano-omics, 7% (17) were identified uniquely by conventional proteomics, and 6% (15) were characterized by both methods. **Fig. 5** depicts the temporal dynamics of a subset, 111 of the 260, of enriched processes and highlights how these processes evolve over time as revealed by the applied analytical approach (conventional vs nano-omics). For the early timepoint samples, response to bacterium, cinnamic acid biosynthesis, defense response by callose deposition in the cell wall, proteolysis, and ROS metabolism were enriched and detectable solely with nano-omics, while cell death, toxin catabolism, and regulation of salicylic acid metabolism were enriched and identified uniquely by conventional proteomics. For the late timepoint samples, programmed cell death, response to endoplasmic reticulum stress, and response to hypoxia were enriched and only resolvable using nano-omics, while systemic acquired resistance, salicylic acid metabolism, and jasmonic acid metabolism were uniquely identified using conventional proteomics. Interestingly, for early timepoint samples, several hormone and metabolite biosyntheses were enriched and had higher gene counts via nano-omics than conventional proteomics, including fatty acid-, glycogen-, jasmonic acid-, lignin-, oxylipin-, phenylacetate- and phosphatidylcholine-biosynthesis. Additionally, proteins involved in quality control for misfolded and/or incompletely synthesized proteins, RuBisCO complex assembly, immune responses, as well as responses to cold, heat, and oxidative stress, were enriched and showed higher gene counts with nano-omics. Glutathione metabolism, response to wounding, and diaminopimelate biosynthesis were also enriched in the early timepoint samples but showed higher gene counts resolvable by conventional proteomics. As a comparative measure, the enriched biological processes associated with the ‘unique’ proteins adsorbed onto the BPEI- (**Supplementary Fig. 12**) and cit-AuNP (**Supplementary Fig. 13**) when incubated with mock samples was also compared. Few proteins associated with responses to cold and heat stressors were slightly enriched on the AuNPs due to the mock treatment. However, responses to oxidative stress or pathogen-induced stress responses were not detected in either AuNP protein corona, suggesting these processes were not upregulated under the mock conditions, as expected. Additionally, as a baseline, the enriched biological processes linked with ‘unique’ proteins from the mock samples relative to the non-infiltrated control were analyzed. **Supplementary Fig. 14** shows that treatment with the mock solution resulted in the overexpression of proteins involved with hyperosmotic salinity responses, protein dephosphorylation, and responses to jasmonic acid stimuli. This finding indicates that infiltration with mock solution slightly alters the proteome of Arabidopsis but does not induce pathogen-dependent stress biomarkers. Collectively, the temporal categorization shown in **Fig. 5** highlights the improved sensitivity and resolution of our nano-omics approach to detecting low-abundance biomarkers of plant stress at early timepoints post-infection.

**Fig. 5.**
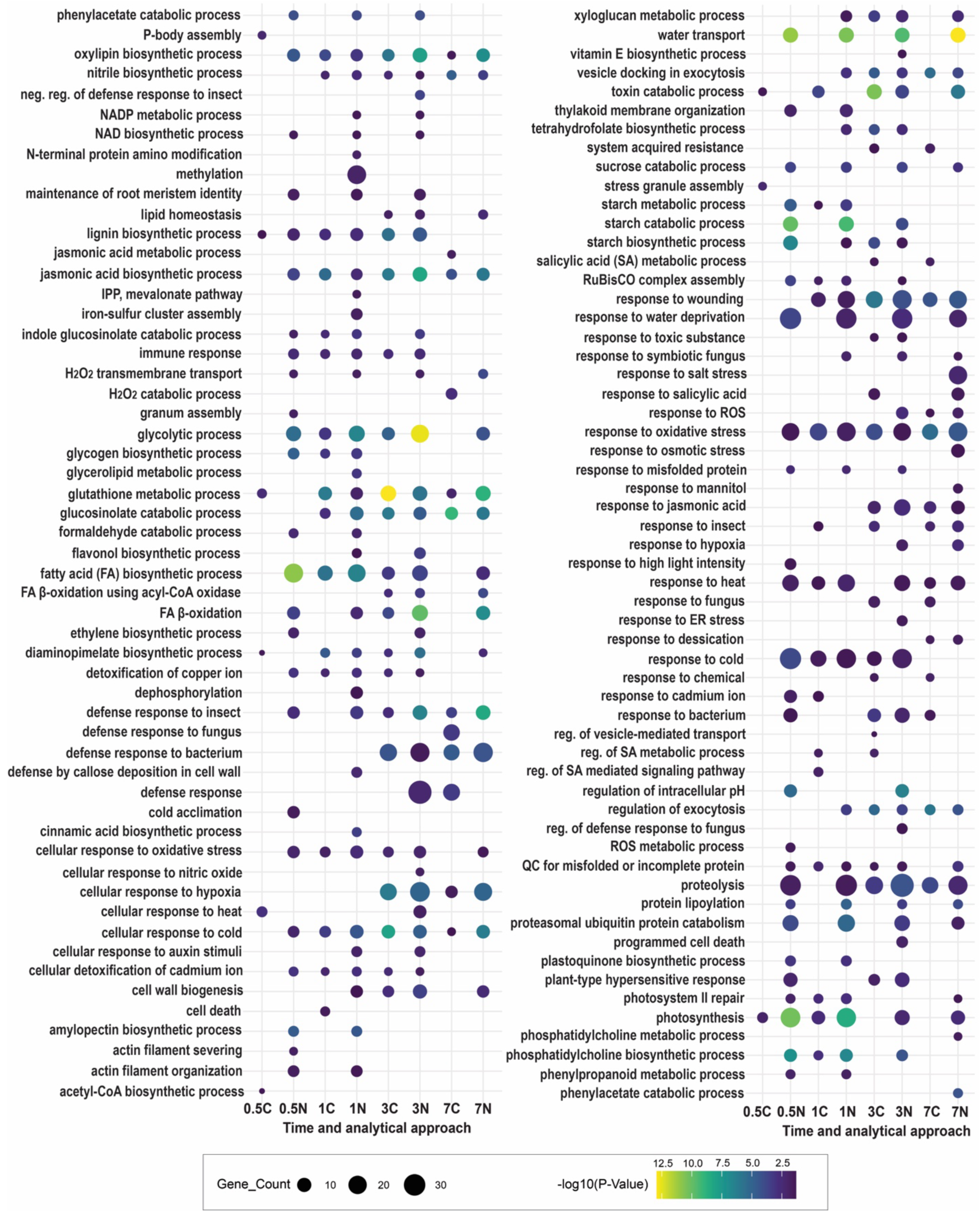
Gene ontology analysis of a subset of enriched biological processes associated with unique proteins identified in pathogen infiltrated lysates and the corresponding AuNP coronas. The x-axis consists of the time-dependent samples analyzed by nano-omics (N) and conventional proteomics (C). The names of the enriched biological processes are displayed on the y-axis. The plot was divided into 2 sections to facilitate better visualization. Gene counts are illustrated by dot size and significance is depicted with a color scale indicative of -log(*p*-values).

In the same manner, gene ontology analysis was conducted on the ‘unique’ proteins identified in distal tissues of pathogen infected plants. A total of 170 enriched biological processes were identified as shown in **Fig. 6**; of these, 55% (94) were detected uniquely by nano-omics, 14% (23) were identified by conventional proteomics, and 31% (53) were discernible by both analytical methods. For the early timepoint distal samples, 54% (91) of the enriched biological processes were identified uniquely by nano-omics, 12% (21) by conventional proteomics, and 21% (36) were detected by both approaches; however, for those that were identified by both, 16% (27) showed higher gene or proteins counts when analyzed by nano-omics. For the late timepoint distal samples, 9% (16) were detected solely by nano-omics, 2% (4) were identified uniquely by conventional proteomics, and 1% (2) were distinguishable by both methods. Notably, for the early timepoint distal samples (0.5- and 1-DPI), water transport, responses to wounding and ROS, methylation, immune response, iron-sulfur cluster assembly, hydrogen peroxide transmembrane transport, and fatty acid β-oxidation were enriched processes that were solely detectable via nano-omics. Contrastingly, only cellular response to heat was enriched and identified uniquely by conventional proteomics in 0.5-DPI distal samples. RuBisCO complex assembly, diaminopimelate and fatty acid biosyntheses were enriched, detected by both analytical approaches, but the associated gene counts were higher via nano-omics. Surprisingly, proteins involved in responses to oxidative stress were expressed early in distal tissues as depicted by their enrichment and analysis via nano-omics, with approximately 20 proteins enriched in 0.5- to 3-DPI distal samples. These proteins were not discernable by conventional proteomics until 7-DPI. For comparison, the enriched biological processes associated with ‘unique’ proteins from distal leaves of mock treated plants were also analyzed (**Supplementary Fig. 15**); the distal leaves showed overexpression of proteins linked with responses to heat and cold, quality control for misfolded of incompletely synthesized proteins, and toxin catabolism. Overall, the findings in **Fig. 6** indicate that there are proteins that are produced in low abundances, at early timepoints, and in tissues distal to pathogen infection that are difficult or not possible to detect with conventional proteomics. However, by applying a nano-omic approach, these low abundant biomarkers of stress, predominately in in distal tissues that were never directly exposed to pathogens, can be enriched and detected more efficiently.

**Fig. 6.**
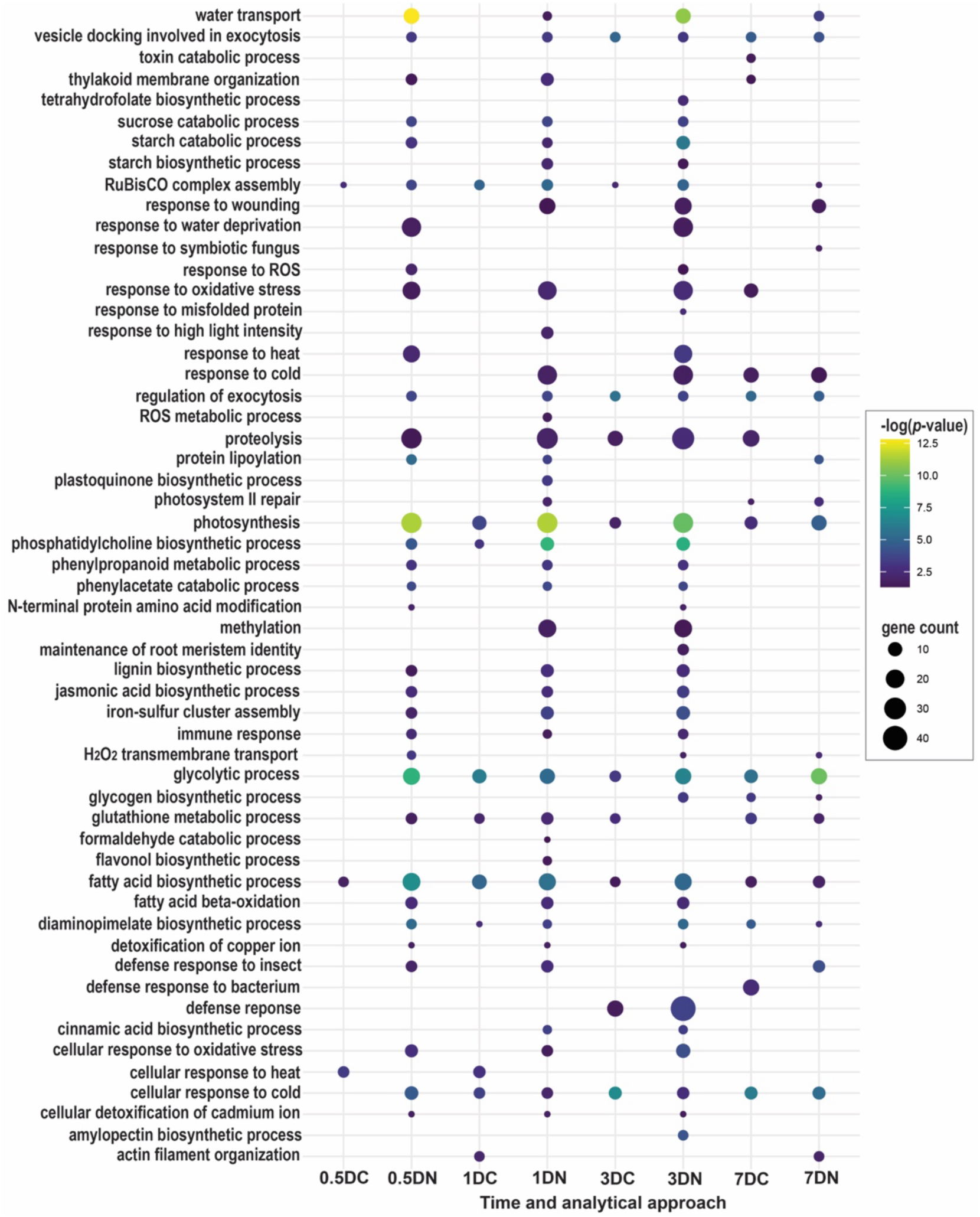
Gene ontology of a subset of enriched biological processes from unique proteins identified in distal tissues collected from pathogen infiltrated plants and their respective AuNP coronas. The x-axis consists of the time-dependent samples analyzed by nano-omics (N) and conventional proteomics (C). Distal tissues are denoted (D) on the x-axis. Names of the enriched biological processes are shown on the y-axis. Gene counts are illustrated by dot size and significance is depicted with a color scale of -log(*p*-values).

Lastly, protein binding affinity, conformation and stability are highly influenced by temperature making it a key factor in determining the composition of nanoparticle protein coronas,^20,41,42^. Consequently, we hypothesized that temperature could potentially impact the reliability and efficiency of our nano-omic approach by altering protein corona compositions. In mammalian studies, protein corona formation is typically performed at 37℃ as this temperature reflects physiological conditions in the human body at which human proteins have evolved to function. However, the optimal temperature for protein corona formation with plant lysates has yet to be established and may depend on specific plant species and their natural environments. We therefore studied temperature dependence of protein corona formation in 3-DPI pathogen infected plant lysates, with AuNP incubated with leaf lysates at both 37℃ and at ambient temperature (∼25℃) because for *A. thaliana* 22–25℃ reflects the conditions at which *in planta* proteins function normally and suppress thermomorphogenesis.^43^ We generally find that while the number of differentially expressed and ‘unique’ *A.thaliana* proteins adsorbed onto the cit- and BPEI-AuNPs were not substantially mediated by temperature, the *P. syringae* proteins identified on the AuNP coronas were indeed temperature dependent. Specifically, ∼2.4 fold more *P. syringae* proteins were characterizable on the AuNP at 37℃ compared to 25℃, (**Supplementary Fig. 16**) highlighting a pivotal temperature-based enrichment of bacterial proteins on both AuNP. Furthermore, as shown in **Supplementary Fig. 17,** the higher temperature resulted in an increased number of adsorbed differentially expressed proteins on the AuNP. However, while 37℃ was optimal to enrich a large quantity of protein, gene ontology analysis indicated that 25℃ was more favorable to deduce significantly impacted biological processes associated with the ‘unique’ proteins from the coronas. **Supplementary Fig. 18** demonstrates that while proteins associated with defense responses to bacterium, responses to oxidative stress, and systemic acquired resistance, were enriched on BPEI- and cit-AuNPs when incubated with pathogen infected samples in a non-temperature dependent manner, proteins linked to innate immunity and responses to bacterium were upregulated in distal tissues of pathogen infected plants and were solely identifiable on the AuNP coronas formed at 25℃. Additionally, for the AuNP coronas formed with the pathogen infiltrated samples, proteins involved in responses to singlet oxygen, siRNA processing, protein-RNA complex assembly, and protein dephosphorylation were enriched uniquely at 25℃. These findings suggest that proteins involved in these critical responses may possess higher conformational stability and/or structural changes that support their greater affinity to the AuNPs at ambient temperatures.

## Discussion

In this work, we aimed to detect the early induction of biotic stress-induced proteins and biomarkers in pathogen infected *A. thaliana*. To accomplish this, we used a nano-omics approach in which cit- and BPEI-AuNP were incubated with infiltrated and distal leaf lysates to form biomolecular coronas for subsequent analysis with UHPLC-MS/MS. This nano-omics approach enabled high-resolution identification and quantification of stress-related proteins and biological processes, offering superior and time-dependent analytical depth compared to conventional proteomics, without selectively depleting highly abundant species from the complex leaf lysates. Moreover, when compared to other standard analytical techniques used to assess plant stress and pathogen infection (**Supplementary Fig. 19**), our nano-omics strategy was most sensitive, detecting early biomarkers of stress in pathogen infected samples prior to the manifestation of phenotypic symptoms of disease, as well as in distal tissues of pathogen infected plants – a feat that was not feasible with customary methods.

Previous studies that have analyzed early stages of differential protein expression in *A. thaliana* infected with *P. syringae* utilized 2-D electrophoresis followed by MS to resolve proteins from complex leaf lysates^44–46^. The dynamic range of protein detection with 2-D electrophoresis can be inherently limited, particularly for underrepresented low abundance proteins^47^. This limitation may have contributed to the relatively low number of stress-related proteins identified in these studies. For example, Sghaier-Hammami, *et al.* identified 24 differentially expressed proteins, of which only 7 were involved in stress defenses.^46^ Similarly, Jones, *et al.* identified 52 differentially expressed proteins, and of those 15 were associated with stress responses.^44,45^ In this work, we identified ∼1500 differentially expressed and ‘unique’ *A.thaliana* proteins using our nano-omics approach, achieving a 2- to 3- fold enhancement in resolution of these proteins when compared to a conventional proteomic approach using high resolution UHPLC-MS/MS. For the early timepoint samples (0.5-DPI) analyzed in this study, 210 differentially expressed and ‘unique’ proteins associated with stress-responses were enriched on the AuNP coronas. Of these 210, none were identified in the 0.5-DPI samples analyzed by conventional proteomics (**Supplementary Fig. 20**), showcasing the superior ability of nano-omics to distinguish and quantify protein variations with greater sensitivity. Moreover, we observed a time-dependent increase of proteins involved in stress-responses with 223, 251 and 198 proteins identified by nano-omics in 1-, 3-, and 7-DPI lysates, respectively. Through this nano-omics approach, we also detected proteins involved in responses to oxidative stress and toxic substances in 0.5-DPI distal leaf lysates, a feature that was not detected with the conventional proteomic approach. These findings reinforce the concept that localized pathogen infections can trigger systemic responses throughput the plant, and that nano-omics may be able to not only detect early plant stress but also can serve to identify new and as-of-yet unknown protein biomarkers of plant stress in the future. Particularly, they highlight the upregulation of proteins involved in stress mitigation and defense mechanisms, enabling the plant to proactively address bacterial infections occurring in distant tissues.

Although the complete proteome of *A. thaliana* has yet to be completely characterized,^48,49^ which is attributable to varying protein abundances across complex tissues and the underrepresentation of proteins from low-abundant transcripts, significant progress has been made with advanced proteomic techniques. However, challenges in characterizing the complete proteomes of understudied plants and crops highlights the need for more sensitive and comprehensive methods, such as nano-omics, to capture a larger range of proteins that improve the depth of proteome coverage. Studies have shown that protein abundances often do not correlate with transcriptional levels in plants.^50,51^ RNA-seq analyses of early transcriptional responses *in planta* have shown that plant defense hormone pathways, such as those mediated by salicylic acid, jasmonic acid, and ethylene, contribute redundantly to plant^52^ and bacterial^53^ transcriptional reprogramming. In this work, the nano-omics approach uniquely identified key immune-related proteins, as well as enzymes associated with jasmonic acid and ethylene biosyntheses, at the earliest timepoint examined (0.5-DPI) – proteins that were undetectable using a conventional proteomic analysis due to their low abundances. Consequently, the proteomic profile generated solely from the conventional analysis might appear negatively correlated with transcriptional data simply because this approach did not resolve these early, low-abundance proteins. In contrast, the nano-omics approach provides a more sensitive and comprehensive early-stage post-transcriptomic analysis, potentially offering a closer alignment with the transcriptional reprogramming that occurs during plant immune responses.

Though AuNP were used in this study to enrich a distinct subset of *A. thaliana* proteins, further studies are needed to optimize nano-omics workflows. This includes identifying the most optimal nanoparticle type(s) with physicochemical properties that enhance the detectable resolution and diversity of plant and agricultural biomolecules. Additionally, we observed that while an ambient temperature was slightly less efficient than the typical 37℃ used in mammalian studies for protein adsorption onto AuNP surfaces, it enabled better enrichment of stress-specific low-abundance biomarkers. These results support the idea that warmer temperatures may promote more dynamic nano-bio interactions driven by increased kinetic energy and collisions but may not always favor the selective enrichment of specific target proteins – further research is needed to understand the influence of other environmental factors on the specificity and sensitivity of nano-omic analyses. Variables such as pH, ionic strength, and biomolecular compositions of different plant tissues and fluids must be explored to fully understand their impact on protein corona formation, and by extension, nano-omic pipelines. Such optimization of nano-omic pipelines has the potential to transform plant proteomics, offering deeper insights into molecular and biological processes in plants that can improve upon agricultural practices, bolster crops, and provide new insights into early stress-induced pathways and mechanisms in plants. This research paves the way for more targeted and field-deployable interventions and detection tools that leverage nanotechnology for sustainable agriculture.

## Methods

### Materials

AuNP were purchased from NanoComposix. Qubit Broad Range protein assay was acquired from Thermo Fisher. Flamingo fluorescent stain, 4× Laemmli buffer and 4-20% SDS-PAGE gels were purchased from BioRad.

### Characterization of AuNP physiochemical properties

Dynamic light scattering (DLS), polydispersity index (PDI), and zeta (ζ) potentials were measured with a Nano Zetasizer (Malvern Panalytical). ζ potential measurements were conducted using Smoluchowski approximation and samples were prepared in ultrapure water (pH 6) with 0.1 mM NaCl for conductivity. UV-Vis spectroscopy was conducted with a UV-3600i Plus Spectrophotometer (Shimadzu).

### Plant growth conditions

Wild-type *A. thaliana* (Col-0) seeds were sown in inundated soil (Sunshine Mix #4), stratified for 3 days at 4°C, and then grown in a growth chamber kept at 24°C with a light intensity of 100-150 µmol/m^2^s. The photoperiod was cycled at 8h light/16h dark. Plants were fertilized on a bimonthly basis with 75 ppm N 20-20-20 general-purpose fertilizer and 90 ppm N calcium nitrate fertilizer reconstituted in H_2_O.

### Preparation of bacterial inocula

*P. syringae* pathovar *tomato* (*Pst*) strain, DC3000, was grown from a stock culture onto an NBY media plate supplemented with rifampicin for 4 days at 30°C. A single bacterial colony was suspended in 10 mM MgCl_2_. The OD_600_ was adjusted to 0.1 (∼ 5 × 10^7^ cfu/mL) by adding 10 mM MgCl_2_ as needed. Serial dilutions were performed to obtain a bacterial suspension with an OD_600_ of 0.001 (∼ 5 x 10^5^ cfu/mL) for leaf infiltration.

### Biotic infection of A. thaliana

At approximately 5-6 weeks, plants were infiltrated with *P. syringae* or with sterile 10 mM MgCl_2_ (mock control) via needless syringe following a standardized protocol for primary leaf inoculation.^23^ Briefly, three leaves, separated 130° from each other, per plant, were infiltrated. Following infiltration, plants were returned to the growth chamber for an allotted time; infections persisted for durations of 0.5-, 1-, 3-, and 7-DPI for leaf collection.

### Photosynthetic activity measurements

Chlorophyll fluorescence measurements were collected with a pulsed amplitude modulation fluorometer equipped with leaf clip holder 2035-B (MINI-PAM-II/B, Walz GmbH, Effeltrich, Germany). Actinic lighting was emitted through optic fibers by a blue (470 nm) LED. The photochemical activity parameters of photosystem II (PSII) were measured by applying a 0.5s saturation pulse (intensity 5000 μmol/m^2^・s) on infiltrated and distal leaves after 1h dark adaptation. The maximum quantum efficiency of PSII primary photochemistry (efficiency at which light adsorbed by PSII is used for photosynthesis when all reaction centers are open) was calculated as F_v_/F_m_.

### Bacterial growth assays

For each timepoint assessed, 1 disc (5.5 mm diameter) was perforated from each of the three infected leaves from 4 different plants. The three discs per plant were pooled as 1 sample, homogenized with 0.1 mL of selective NBY broth via bead beating, and subjected to a 1:10 dilution series. The samples (0.1 mL) were plated onto selective NBY plates and incubated at ambient temperature for 2 days before colony forming units (CFU) were counted.

### Protein corona formation and extraction

Leaves from both distal and primary infection sites were collected from plants treated with *P. syringae* as well as those subjected to mock treatments. To eliminate the risk of false positives in distal tissues, potentially caused by epiphytic stages of *P. syringae*,^54^ the collected distal leaves were sequentially submerged in 70% EtOH, 10% NaClO, and ultrapure H_2_O briefly to sterilize foliar surfaces. Leaves were immediately flash-frozen in liquid nitrogen and homogenized in ultrapure H_2_O using a bead beater. Leaf lysates were centrifuged 1h at 5,000 × g (4°C) to pellet chloroplasts and cell wall debris. Lysates were normalized to 0.1 mg/mL after quantifying protein concentrations with the Qubit Broad Range assay. Protein corona was formed with 20 µg/mL AuNPs which were added to leaf lysates (50 µL final volume; final protein concentration 0.1 mg/mL) and incubated at 37°C or ambient temperature for 1h. Following protein corona formation, AuNPs were pelleted (21,000 × g; 30 min) and washed thrice with 1× PBS. Proteins were denatured and desorbed from resuspended AuNPs (10 μL, 1× PBS) with (1:1, v/v) 4× Laemmli buffer. For the time-dependent nano-omics approach, proteins were separated using 4% SDS-PAGE (110 V; 10 min). For the temperature-dependent nano-omics approach, proteins were separated using 4-20% SDS-PAGE (110 V; 65 min). Gels were stained with 1× Flamingo fluorescent gel stain solutions and imaged with a GE Typhoon FLA 9000 Imaging Scanner.

### In gel digestion of proteins

For gels that were run for 10 mins, the proteins did not separate into individual bands and the singular band was excised for downstream analysis. For gels that were run to completion (1h), protein bands (∼ 75-25 kDa) were excised from the SDS-PAGE gels. Gel bands were then diced into 1mm × 1 mm cubes and unstained 3× by first washing with 100 μL of 100 mM ammonium bicarbonate (NH_4_HCO_3_) for 15 min followed by an addition of 100 μL of acetonitrile for 15 min. The supernatants were removed, and the gel pieces were dried with a SpeedVac. Samples were reduced by incubating the gel pieces with 200 μL of 10 mM DTT in 100 mM NH_4_HCO_3_ at 56°C for 30 min. After cooling to room temperature, the supernatants were removed and replaced with 200 μL of 55 mM IAA in 100 mM NH_4_HCO_3_ and the samples were incubated at room temperature in the dark for 20 min. The supernatants were then removed, and the gel pieces were washed 1× with 200 μL of 100 mM NH_4_HCO_3_ for 15 min. Samples were then dehydrated with 200 μL acetonitrile and dried with a SpeedVac. For protein digestion, enough solution of ice-cold trypsin (0.01 μg/μL), in 50 mM NH_4_HCO_3_, was added to cover the gel pieces and the samples were placed on ice for 30 min. After complete rehydration of the gel pieces, the trypsin solutions were removed and replaced with 50 mM NH_4_HCO_3_ and left overnight at 37°C. The peptides were extracted 2× by adding 50 μL of 0.2% formic acid/ 5% acetonitrile and vortexing the samples for 30 min at ambient temperature. The supernatants containing peptides were collected. Any remaining peptides were further extracted from the gel pieces by adding 50 μL of 0.2% formic acid/ 50% acetonitrile and vortexing for 30 min at ambient temperature. The supernatants were collected, pooled with the peptides from the first extraction, and dried.

### UHPLC-MS/MS

Dried peptides were redissolved in 10 μL of 5% formic acid and analyzed by UPHLC-MS/MS using nano-electrospray ionization (nano-ESI). The nano-ESI was performed using a timsTOF Pro 2 hybrid mass spectrometer (Bruker) interfaced with nano-scale reversed phase UHPLC (EVOSEP ONE). Mobile phase A was comprised of 0.1% formic acid and mobile phase B was composed of 0.1% formic acid/ 99.9% acetonitrile. The timsTOF Pro 2 MS was operated in the PASEF mode for standard proteomics. Protein identification and label free quantification was carried out using Peaks Studio X (Bioinformatics Solutions, Inc) against the UniProt *A. thaliana* and *P. syringae* databases (UP000006548 & UP000002515, respectively).

### Gene ontology analyses

UniProt accession IDs were input into the Functional Annotation Tool of the DAVID bioinformatics platform for gene ontology analysis. For this analysis, thresholds were set to a count of 2 and an EASE value of 0.05 for the identification of enriched biological processes associated with input list of protein uniport accession IDs. UniProt accession IDs were also input into the STRING database for gene ontology analysis dependent on protein-protein interaction networks. For this analysis, a maximum false discovery rate of ≤ 0.05 was used to analyze enriched biological processes.

### Statistics and reproducibility

Statistical analysis and visualization were performed with GraphPad Prism (v.10.1.1) and R (v.4.3.2), respectively. The following packages were used in R to conduct PCA, generate hierarchical heatmaps, and create dots plots in R: stats, factoextra, pheatmap, and ggplot2. Experiments were conducted with biological triplicates (*n* = 3), unless otherwise noted.

## Acknowledgements

R.C. is supported by the NSF PRFB award number 2305663 and the Burroughs Wellcome Fund (BWF) PDEP award. H.J.S. and E.V. are supported by the DOD through the NDSEG graduate fellowship. We further acknowledge support of a BWF Career Award at the Scientific Interface (CASI) (to M.P.L.), a Dreandyfus foundation award (to M.P.L.), the Philomathia foundation (to M.P.L.), an NIH MIRA award (to M.P.L.), an NIH R03 award (to M.P.L), an NSF CAREER award (to M.P.L), an NSF CBET award (to M.P.L.), an NSF CGEM award (to M.P.L.), a CZI imaging award (to M.P.L), a Sloan Foundation Award (to M.P.L.), a USDA BBT EAGER award (to M.P.L), a Moore Foundation Award (to M.P.L.), an NSF CAREER Award (to M.P.L), and a DOE Office of Science grant with award number DE-SC0020366 (to M.P.L.). M.P.L. is a Chan Zuckerberg Biohub investigator, a Hellen Wills Neuroscience Institute Investigator, and an IGI Investigator.

## Author Contributions

R.C. conceived and designed the project. R.C. carried out the experiments for this project with help from N.S, H.J.S., and E.V. Data analysis was conducted by R.C. with support from T-J. L. The manuscript was written by R.C. Technical guidance, research prioritization, manuscript writing and editing was provided by M.P.L. All authors provided editorial guidance on the manuscript.

## Ethics

Due diligence was taken to ensure all co-authors were properly credited for their contributions to this work.

