## Supplementary Information for "Deep profiling of plant stress biomarkers following bacterial pathogen infection with protein corona based nano-omics"

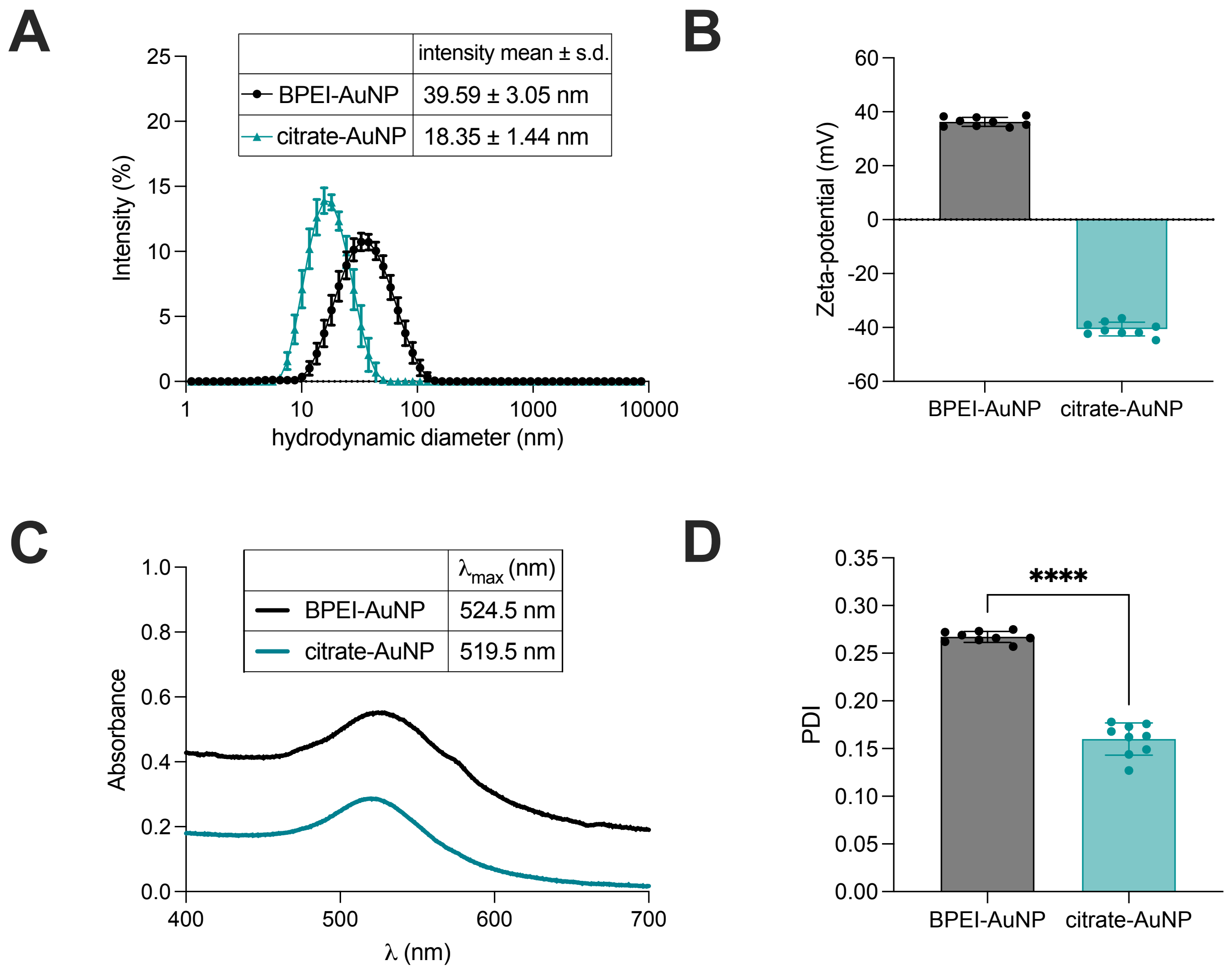


**Supplementary Fig. 1.** Characterization of AuNPs. **A)** Intensity means ± standard deviations of the hydrodynamic diameters of BPEI conjugated and citrate capped AuNPs were measured with DLS ($n$=9, 3 technical replicates per sample replicate). **B)** $\zeta$-potential measurements of the AuNPs. ($n$=9) **C)** UV-vis spectra of the AuNPs used in this study. **D)** PDI values for the AuNPs. ($n$=9). All measurements were conducted with AuNPs at a final concentration of 20 µg/mL. An unpaired $t$-test was conducted for PDI measurements; asterisks represent statistical significance with **** indicating $p$-values < 0.0001.

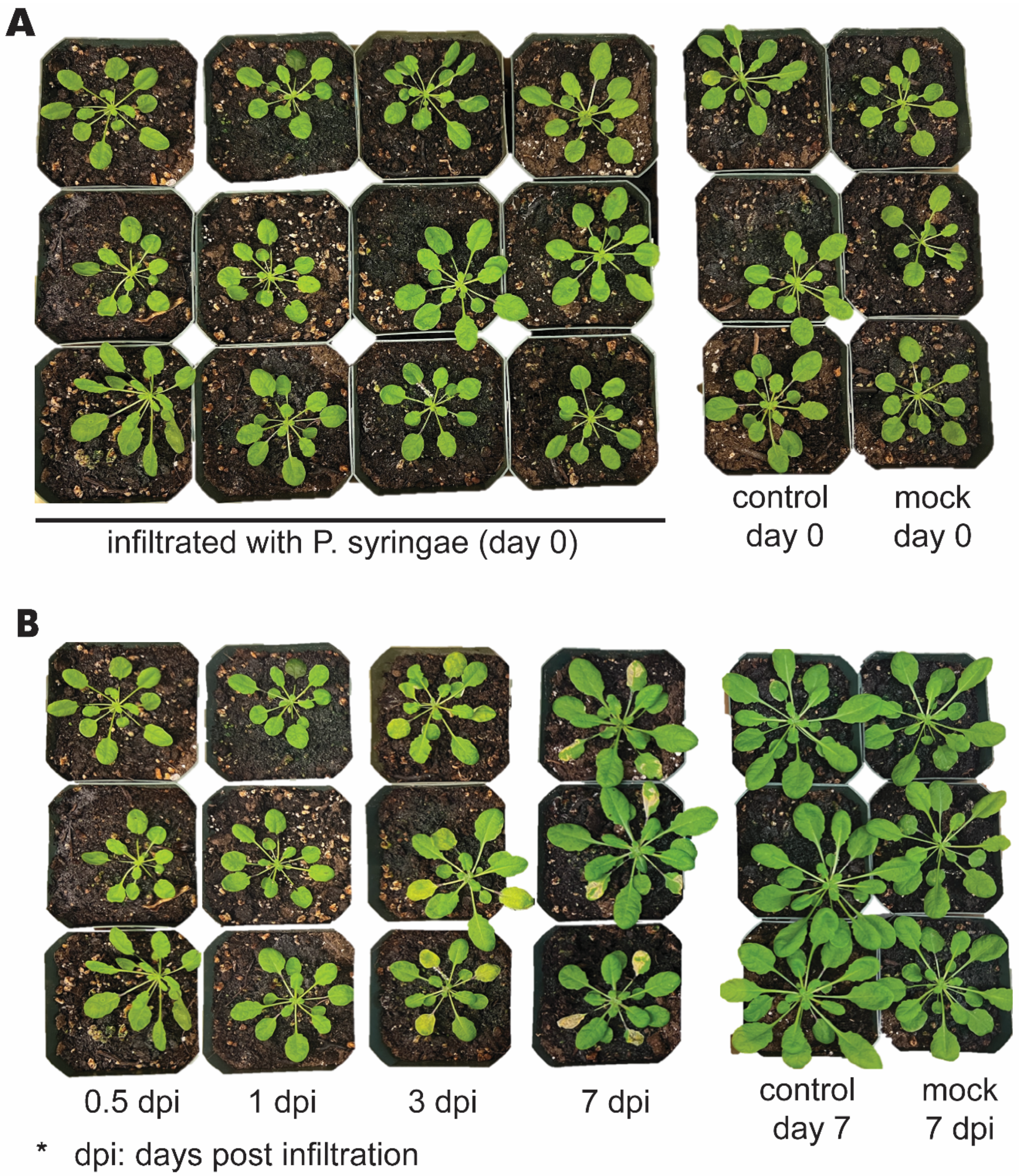


**Supplementary Fig. 2.** Photo images of *A. thaliana* **A)** Five-week-old plants were infiltrated with P. syringae or mock solution on day 0. **B)** Infected plants were photographed and collected for proteomics 0.5, 1, 3 and 7 days following the pathogen infiltration. Mock treated plants and untreated (control) plants were photographed and collected 7 days after day 0.

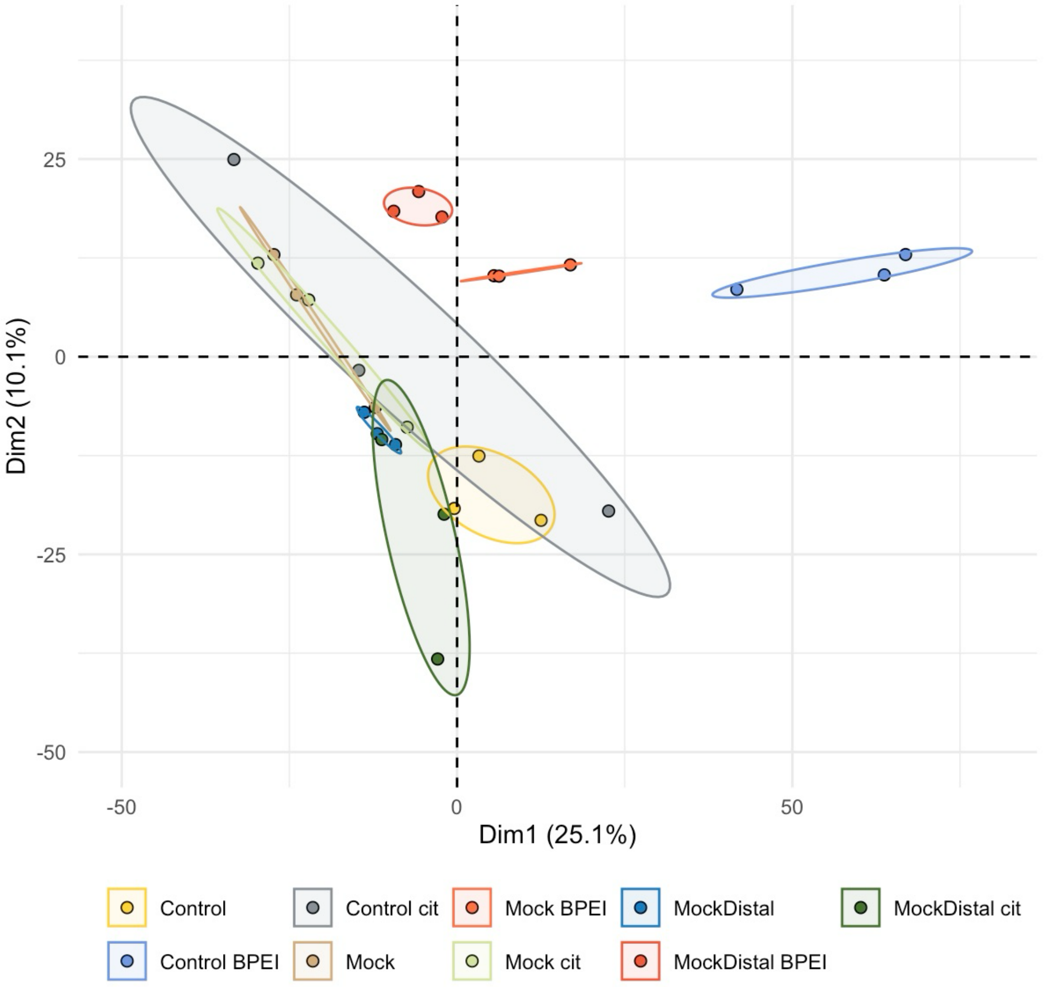


**Supplementary Fig. 3.** Score plot of the PCA conducted with the protein spectral counts in the mock, mock distal, non-infiltrated control, and their respective cit-AuNP and BPEI-AuNP coronas. Each point represents a biological replicate, and ellipses represent the 95% confidence intervals around the mean point of each group (*n*=3). PC1 and PC2 summarized 35.2% of the variance.

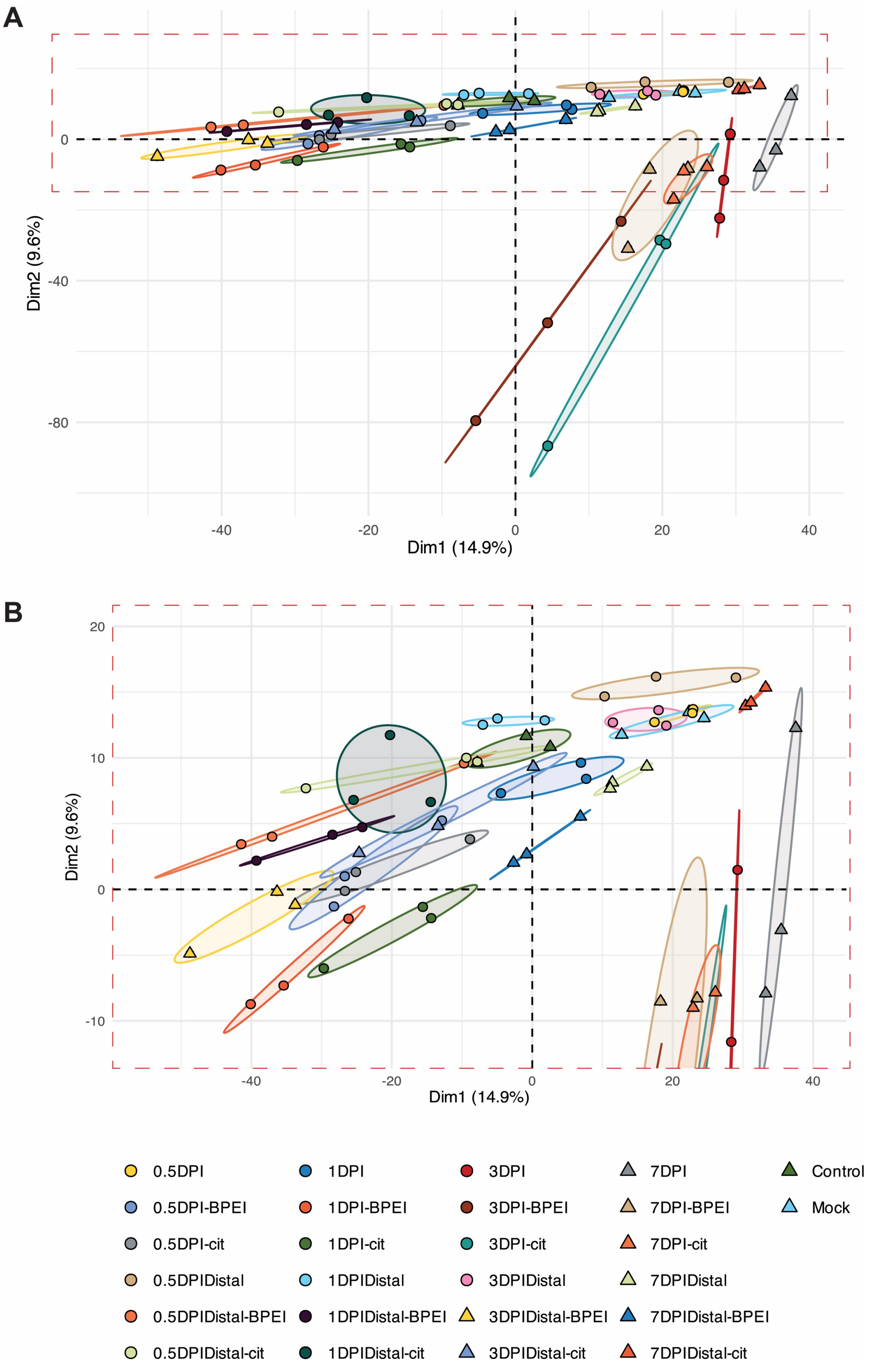


**Supplementary Fig. 4. A)** Score plot of the PCA conducted with protein spectral counts with all samples. Each point represents a biological replicate, and ellipses represent the 95% confidence intervals around the mean point of each group (*n*=3). PC1 and PC2 summarized 24.5% of the variance. **B)** Magnified view of the samples within the red dashed box in the PCA plot (panel A).

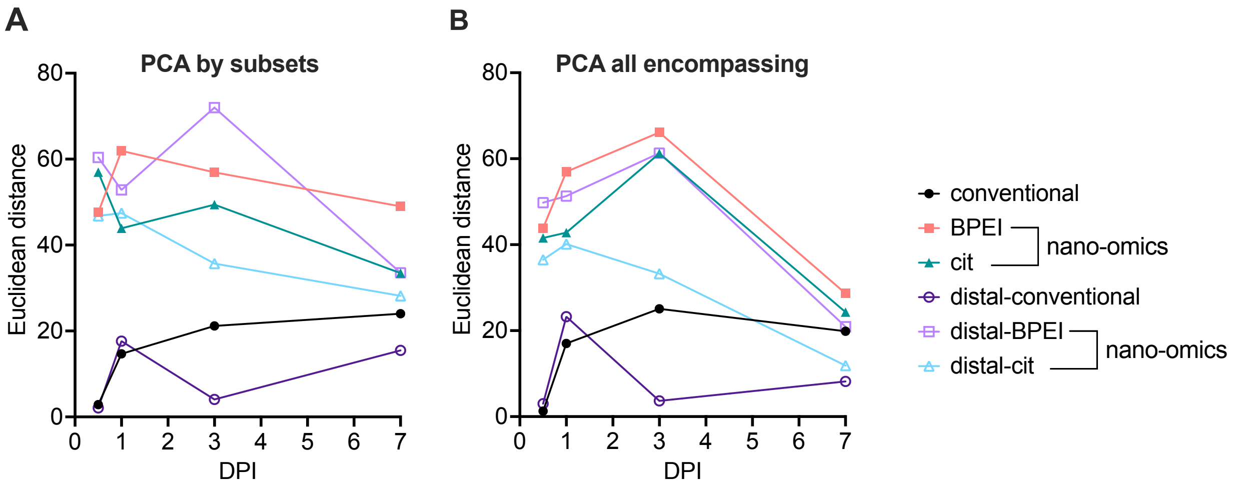


C

| | | nano-omics | | | nano-omics | |
|---|---|---|---|---|---|---|
| DPI | conventional | BPEI | cit | distal-conventional | distal-BPEI | distal-cit |
| 0.5 | 2.867844969 | 47.58143804 | 56.972523930 | 2.097496876 | 60.40259581 | 46.782219940 |
| 1.0 | 14.693002900 | 61.92228451 | 43.920871530 | 17.621786310 | 52.84739079 | 47.443454350 |
| 3.0 | 21.172950200 | 56.92739782 | 49.422035780 | 4.055972528 | 71.98167605 | 35.687208810 |
| 7.0 | 24.006126790 | 49.02754772 | 33.459468660 | 15.478703740 | 33.53848356 | 28.167788140 |

D

| | | nano-omics | | | nano-omics | |
|---|---|---|---|---|---|---|
| DPI | conventional | BPEI | cit | distal-conventional | distal-BPEI | distal-cit |
| 0.5 | 1.306407301 | 43.88209643 | 41.573892010 | 3.021739441 | 49.76024491 | 36.507094940 |
| 1.0 | 17.014738650 | 56.94623063 | 42.807431000 | 23.247168980 | 51.29766374 | 40.139382800 |
| 3.0 | 25.090613120 | 66.11732411 | 61.215954200 | 3.661533060 | 61.29251859 | 33.262501600 |
| 7.0 | 19.858515800 | 28.66347546 | 24.299203190 | 8.175909808 | 20.91696792 | 11.871585180 |

**Supplementary Fig. 5.** Euclidean distances from the mock, calculated using the average PCA coordinates of each sample type, plotted over time. **A)** shows distances calculated using PCAs conducted on subsets of samples (see **Fig. 2**), while **B)** illustrates the distances calculated using a PCA that included all samples during the analysis. Tables in panels **C)** and **D)** list the Euclidean distance values, respectively.

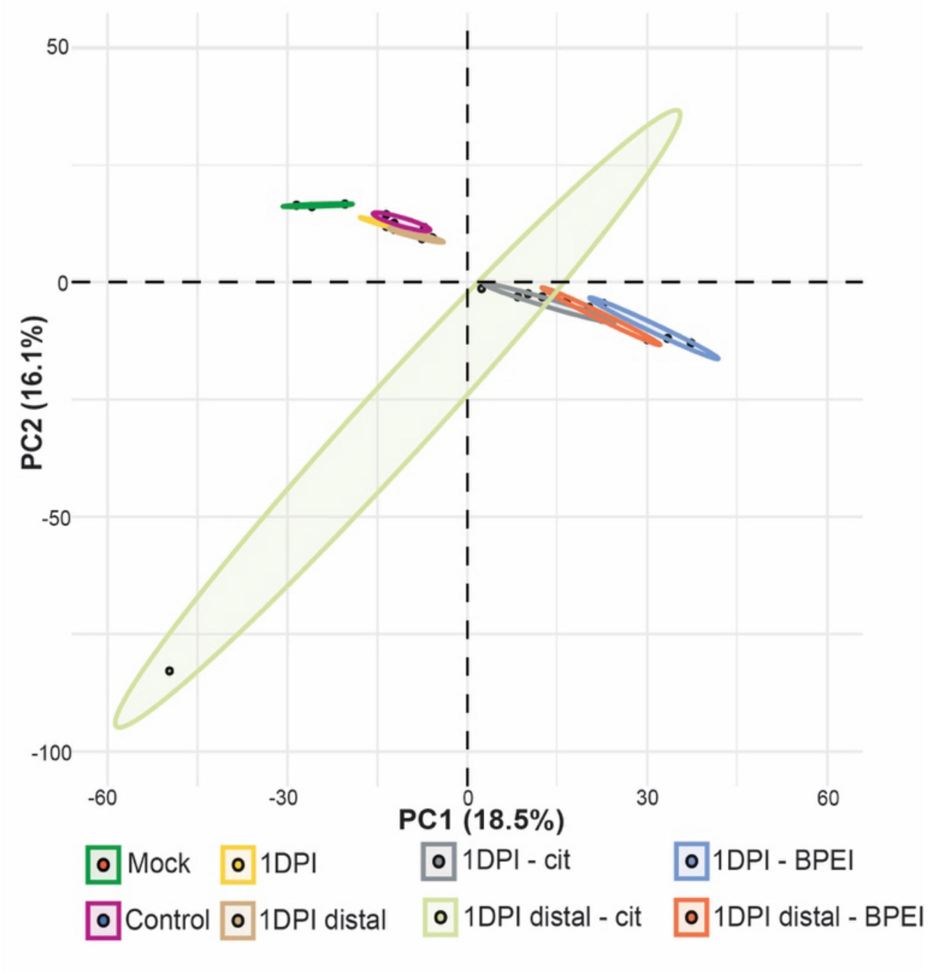


**Supplementary Fig. 6.** Score plot of the PCA conducted with the protein spectral counts in the mock, non-infiltrated control, 1-DPI and 1-DPI distal lysates, and their respective coronas. Each point represents a biological replicate, and ellipses represent the 95% confidence intervals around the mean point of each group (*n*=3). PC1 and PC2 summarized 34.6% of the variance.

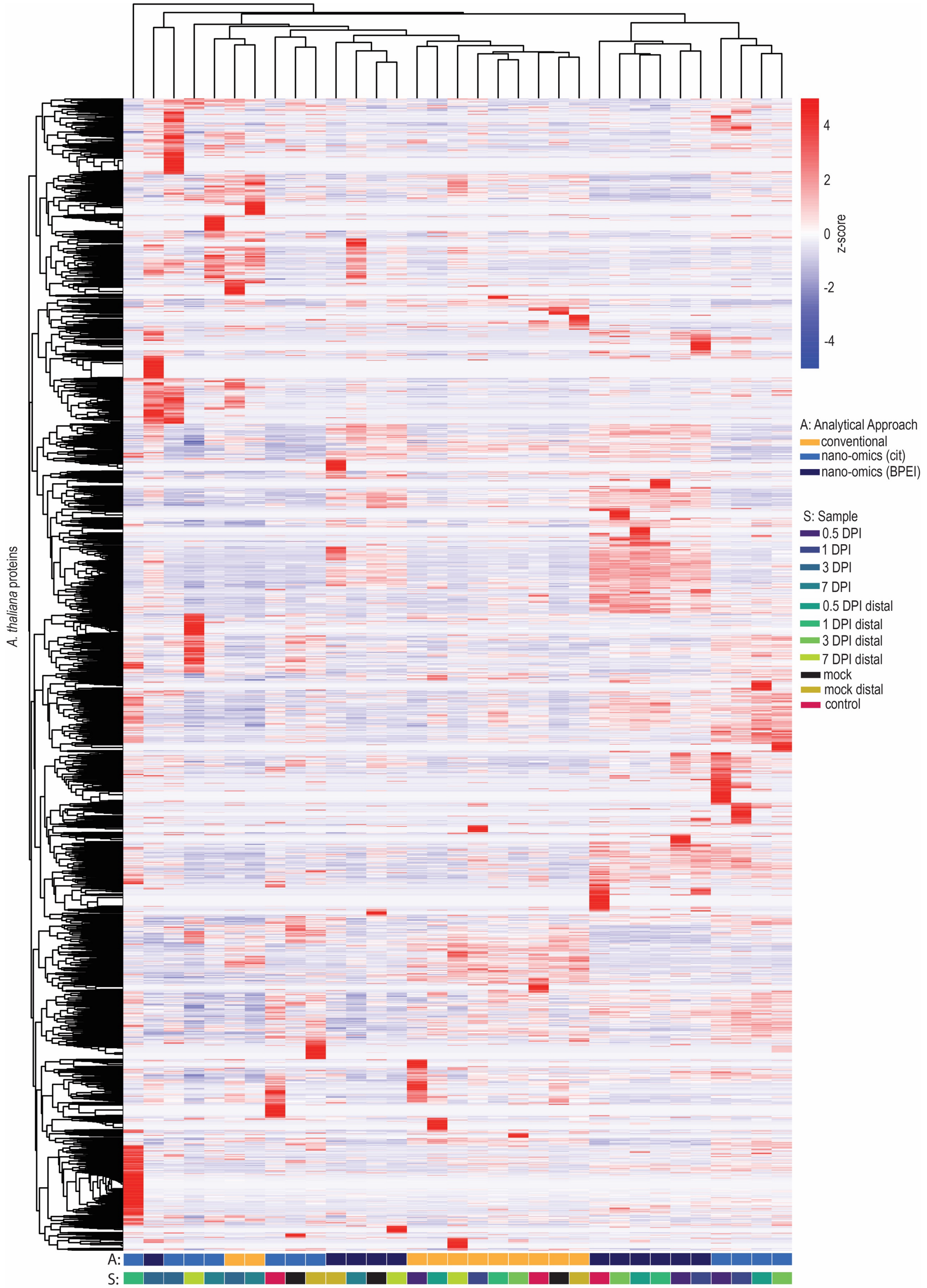


**Supplementary Fig. 7.** Heatmap with hierarchical clustering (Euclidean, with a complete clustering method using the pheatmap ackage in R) of z-scores calculated from the average relative abundance of *A. thaliana* proteins. The heatmap scale depicts z-scores. The y-axis represents proteins and the x-axis includes the sample types (labeled as 'S') analyzed by different analytical approaches (labeled as 'A').

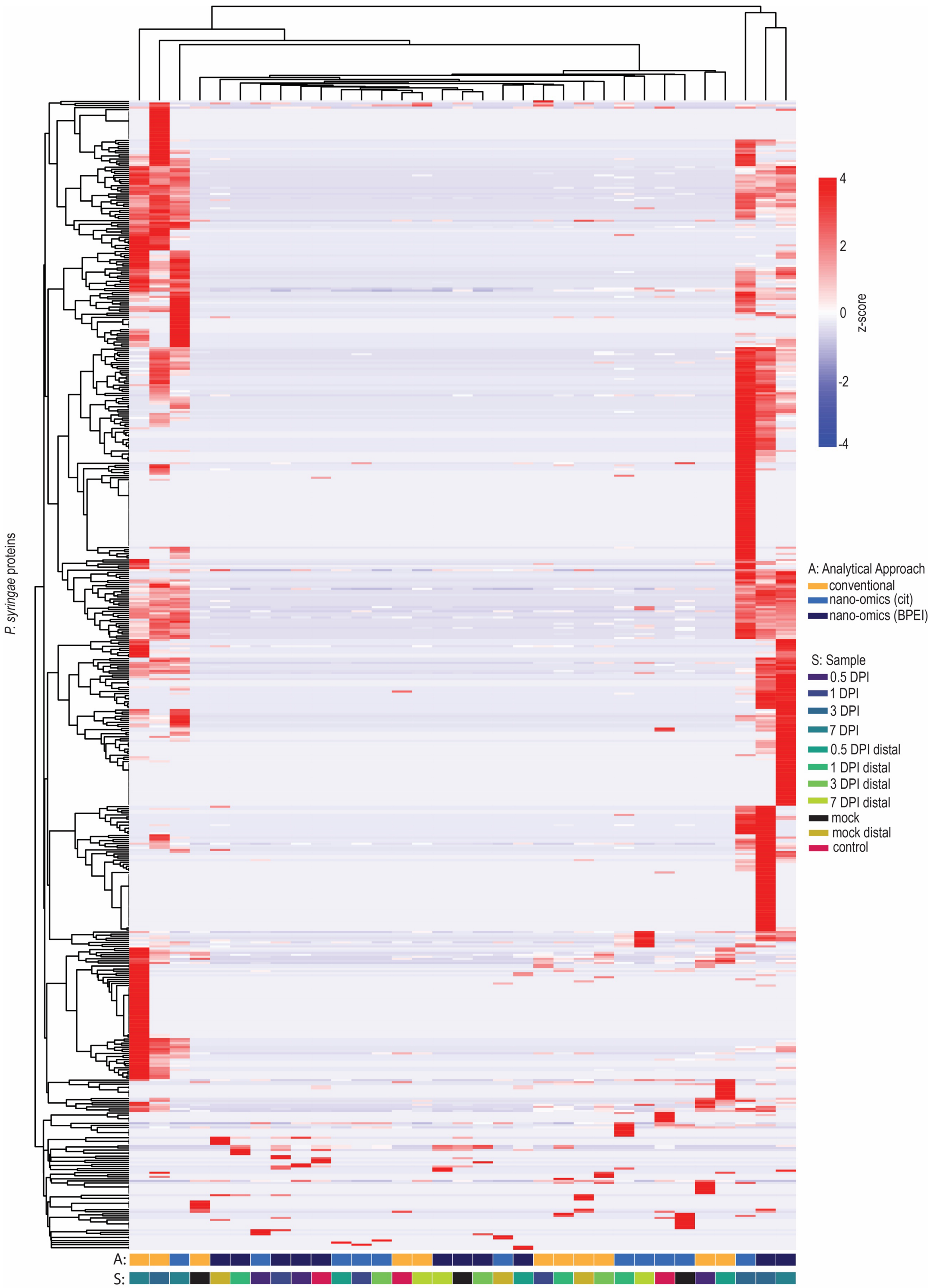


**Supplementary Fig. 8.** Heatmap with hierarchical clustering (Euclidean, with a complete clustering method using the pheatmap package in R) of z-scores calculated from the average relative abundance of *P. syringae* proteins in each sample. The heatmap scale denotes z-scores. The y-axis represents proteins and the x-axis includes the sample types (labeled as 'S') analyzed by different analytical approaches (labeled as 'A').

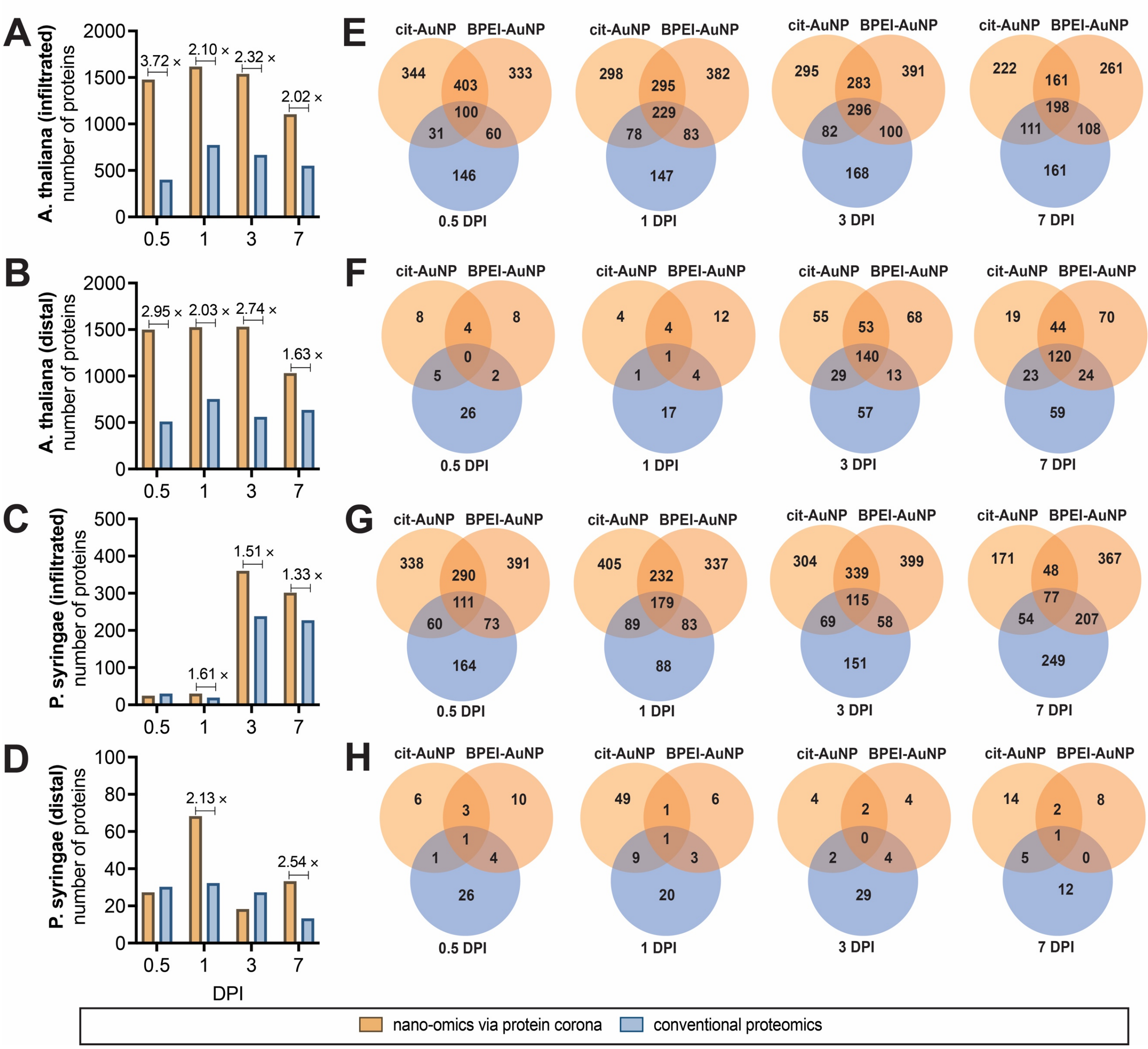


**Supplementary Fig. 9.** Quantity of differentially expressed proteins [log2(fc) >1 or <-1] and unique proteins, identified relative to protein expression in mock treated plants, by conventional proteomic (blue) and nano-omic (orange) approaches. The number of differentially expressed and unique *A.thaliana* and *P. syringae* proteins identified in **A), C)** pathogen infiltrated *A.thaliana* leaves and **B), D)** distal leaves of pathogen infiltrated *A.thaliana* plants, respectively. For the nano-omic bars, the quantity of proteins from both AuNP coronas were combined. Venn diagrams illustrate the commonalities and differences between the quantity of differentially expressed and unique **E), F)** *A. thaliana* and **G), H)** *P. syringae* proteins identified by the nano-omic and the conventional proteomic approach.

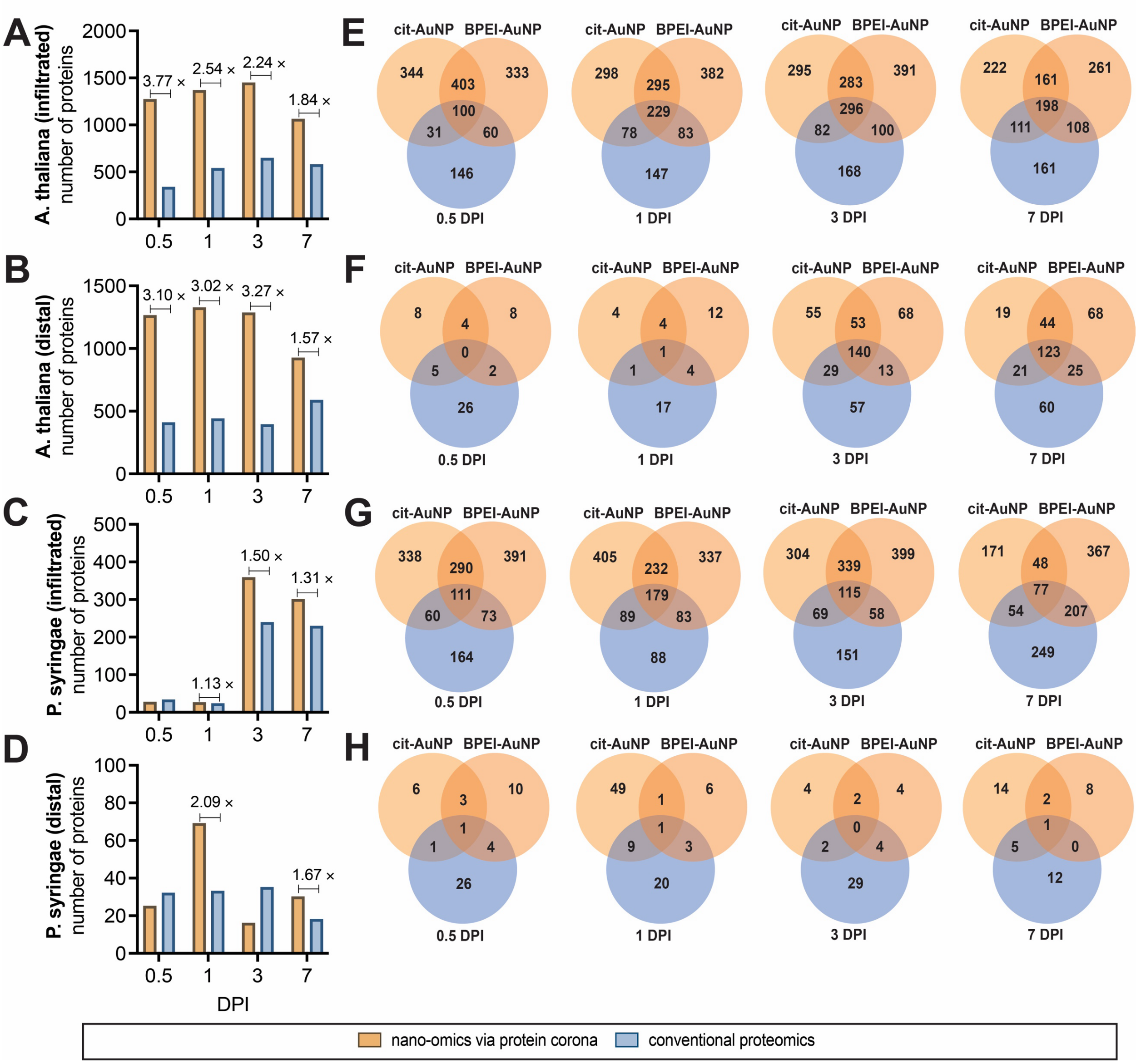


**Supplementary Fig. 10.** Quantity of differentially expressed proteins [log2(fc) >1 or <-1] and unique proteins, identified relative to protein expression in non-infiltrated controls, by conventional proteomic (blue) and nano-omic (orange) approaches. The number of differentially expressed and unique *A.thaliana* and *P. syringae* proteins identified in **A), C)** pathogen infiltrated *A.thaliana* leaves and **B), D)** distal leaves of pathogen infiltrated *A.thaliana* plants, respectively. For the nano-omic bars, the quantity of proteins from both AuNP coronas were combined. Venn diagrams illustrate the commonalities and differences between the quantity of differentially expressed and unique **E), F)** *A. thaliana* and **G), H)** *P. syringae* proteins identified by the nano-omic and the conventional proteomic approach.

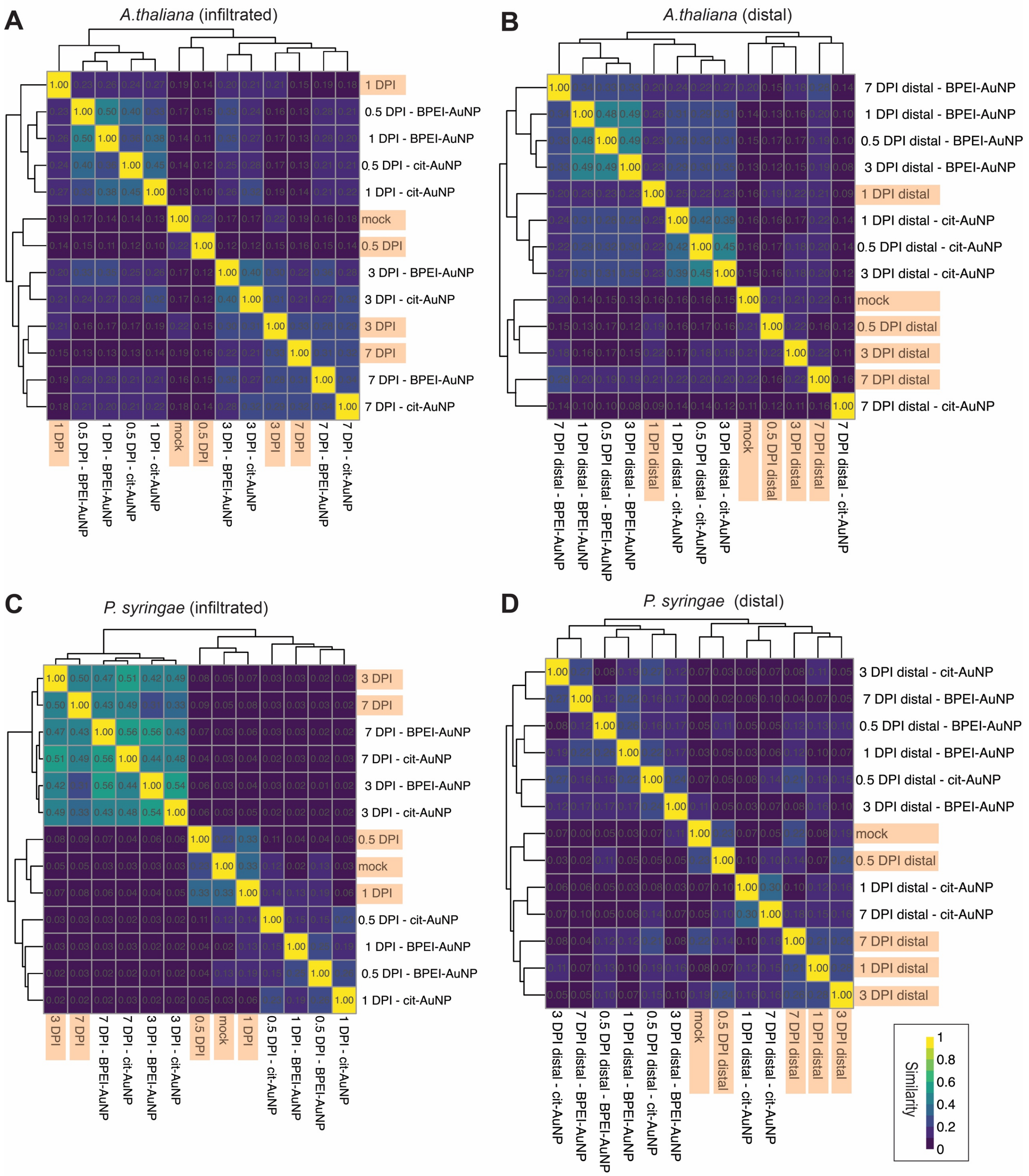


**Supplementary Fig. 11.** Heatmaps with hierarchical clustering depict the similarities between samples based on the composition of **A), B)** *A. thaliana* and **C), D)** *P. syringae* differentially expressed ($\log_2$(fc)>1 and <-1) and unique proteins, relative to non-infiltrated controls, identified in **A), C)** pathogen infiltrated and **B), D)** distal tissues of pathogen infected plants. The color scale indicates the degree of similarity based on Jaccard index values. Samples labeled in orange represent those analyzed by the conventional proteomic approach.

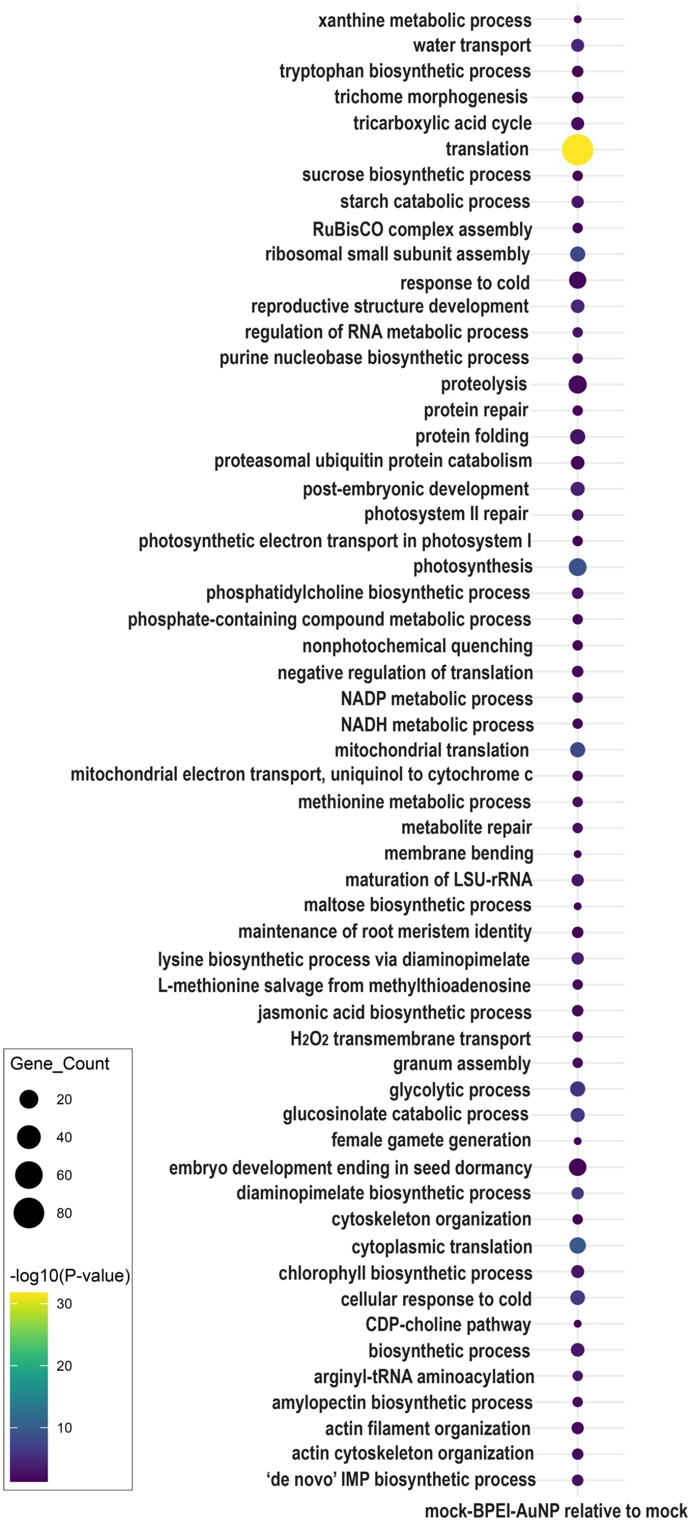


**Supplementary Fig. 12.** Gene ontology analysis of enriched biological processes from unique proteins detected on the BPEI-AuNP corona formed with the mock samples (analyzed by the nano-omic approach) relative to the protein expression in the mock samples (analyzed by the conventional proteomic method). Gene counts are illustrated by dot size and significance is depicted with a color scale of -log(*p*-values).

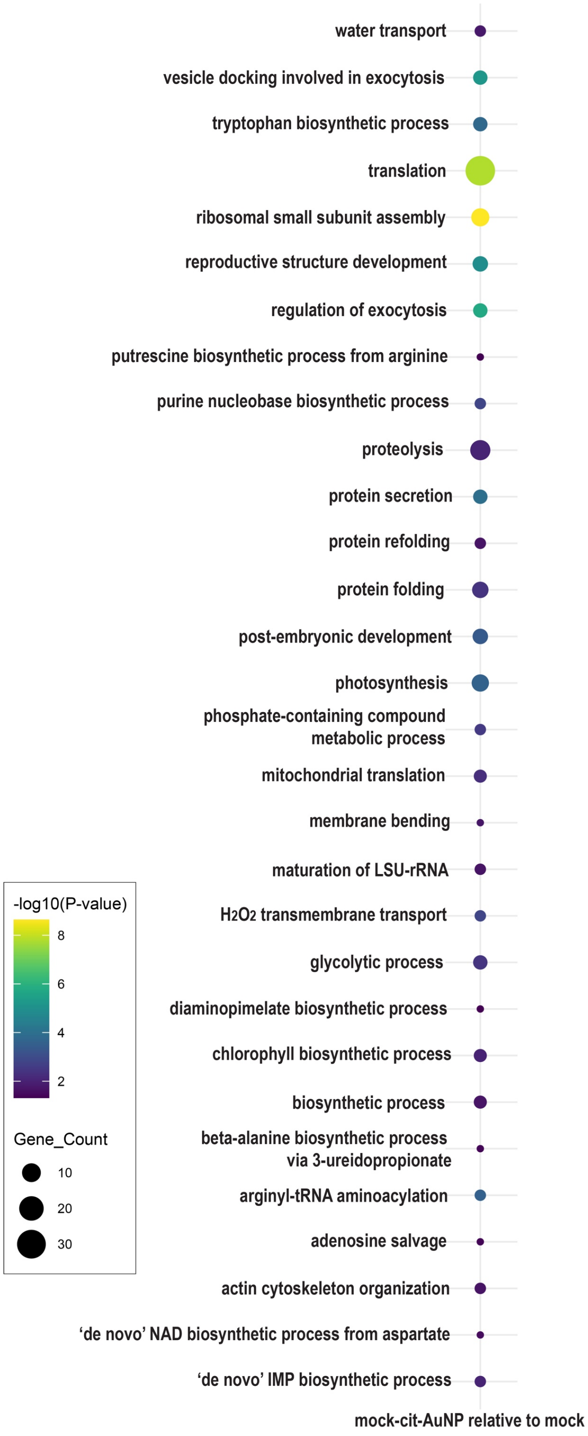


**Supplementary Fig. 13.** Gene ontology analysis of enriched biological processes from unique proteins detected on the cit-AuNP corona formed with the mock samples (analyzed by the nano-omic approach) relative to the protein expression in the mock samples (analyzed by the conventional proteomic method). Gene counts are illustrated by dot size and significance is depicted with a color scale of -log(*p*-values).

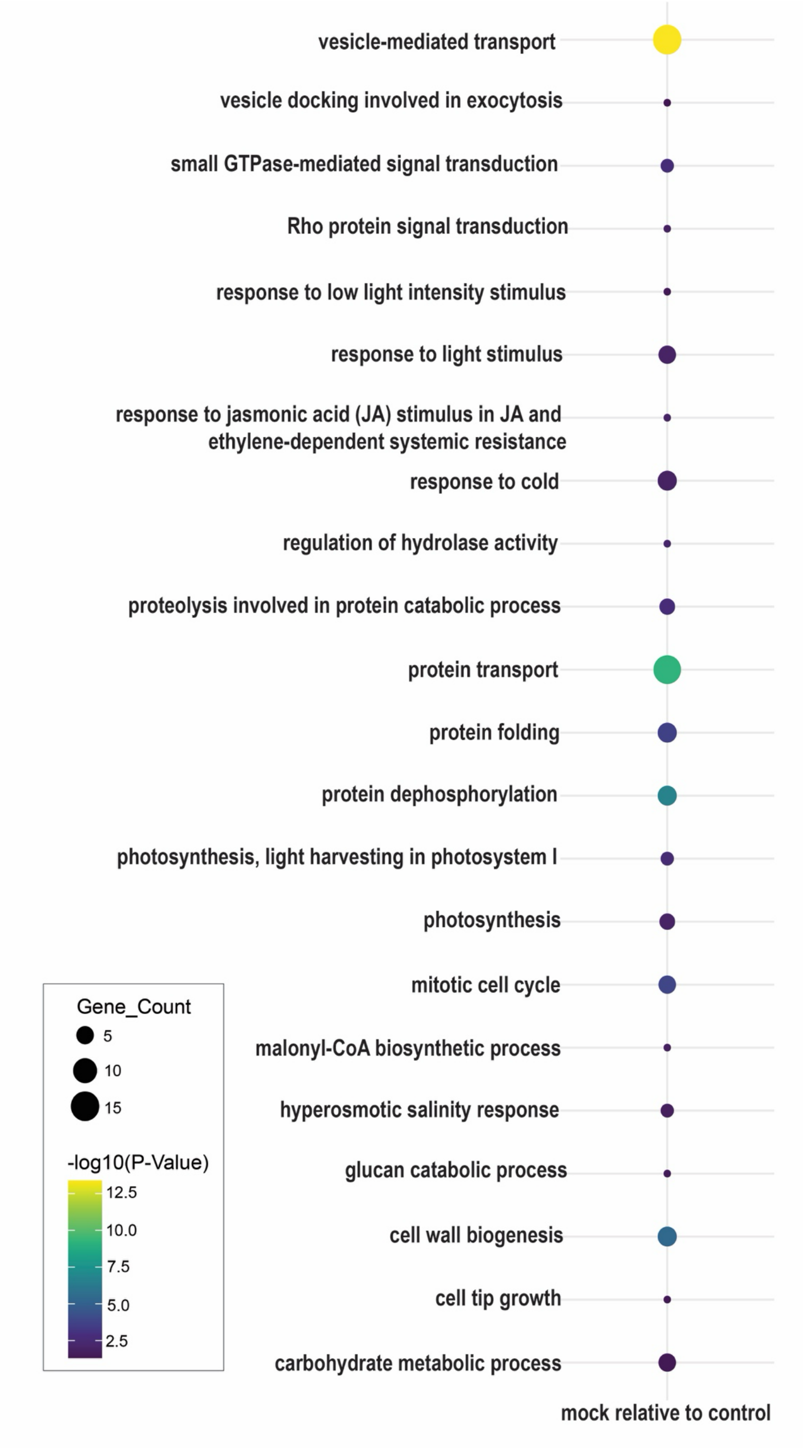


**Supplementary Fig. 14.** Gene ontology analysis of enriched biological processes from unique proteins identified in the mock samples (analyzed by the conventional proteomic method) relative to the protein expression in the non-infiltrated control samples (analyzed by the conventional proteomic method). Gene counts are illustrated by dot size and significance is depicted with a color scale of -log(*p*-values).

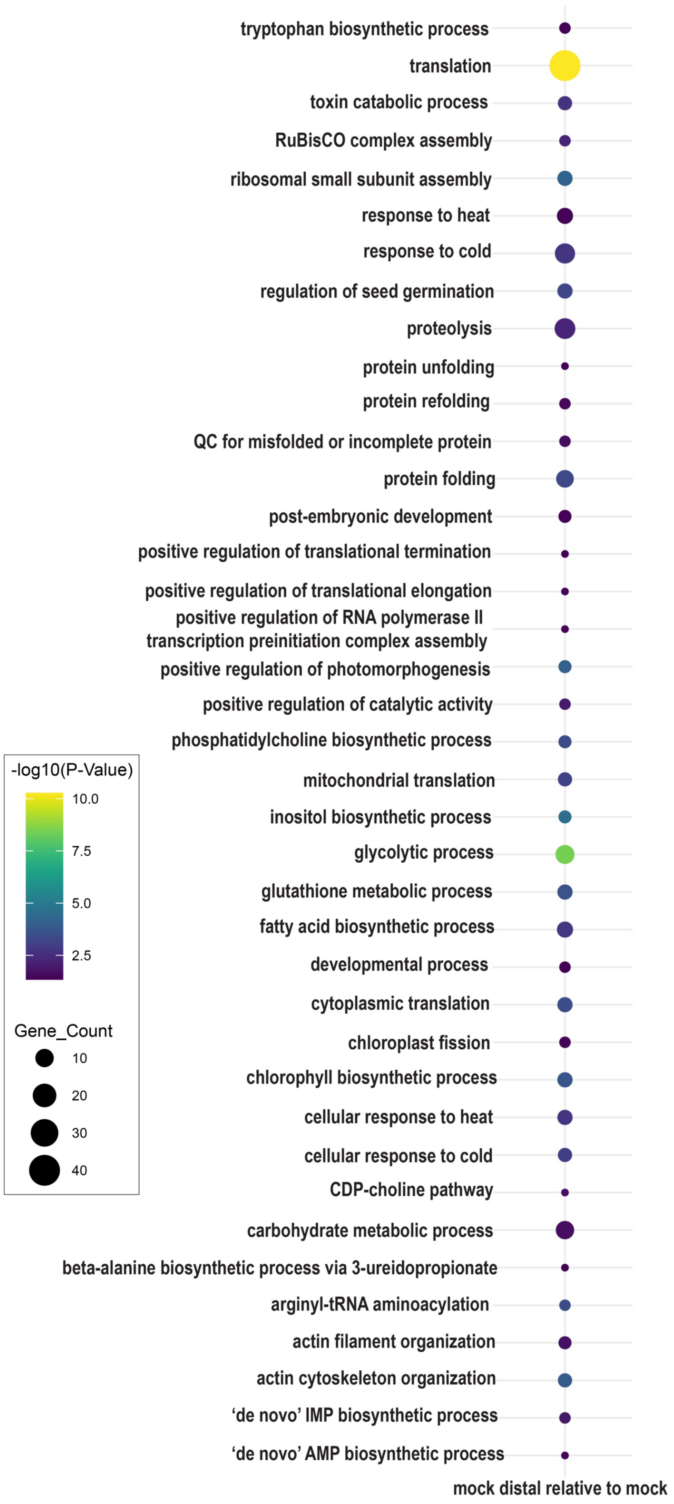


**Supplementary Fig. 15.** Gene ontology analysis of enriched biological processes from unique proteins identified in the mock distal samples (analyzed by the conventional proteomic method) relative to the protein expression in the mock treated samples (analyzed by the conventional proteomic method). Gene counts are illustrated by dot size and significance is depicted with a color scale of -log($p$-values).

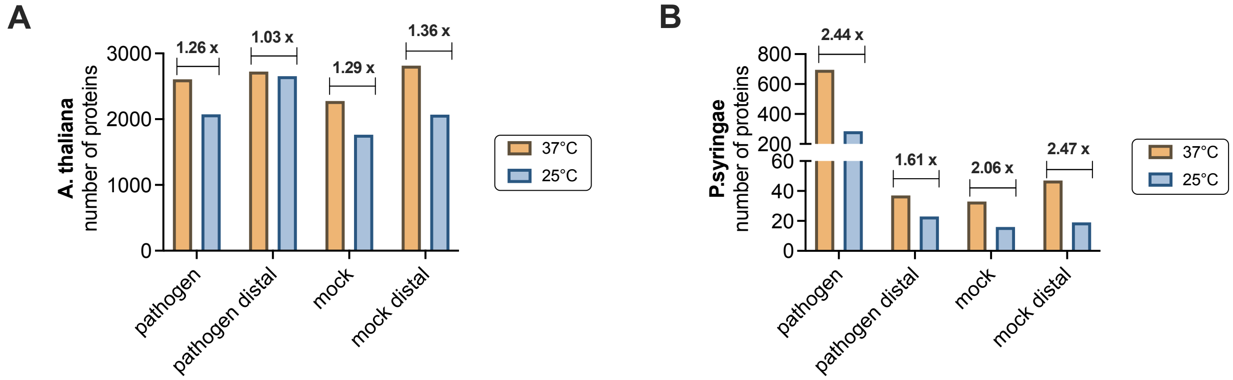


**Supplementary Fig. 16.** Temperature dependence on the quantity of differentially enriched (log2(fc)>1 and <-1) and 'unique' **A)** *A. thaliana* and **B)** *P.syringae* proteins analyzed from both AuNP coronas. The coronas were formed in 3-DPI pathogen infiltrated plants, mock infiltrated plants, and their respective distal leaf lysates at 37°C (orange bars) and 25°C (light blue bars). Protein expression on the coronas (analyzed by nano-omics) was compared relative to mock treated plants (analyzed by conventional proteomics).

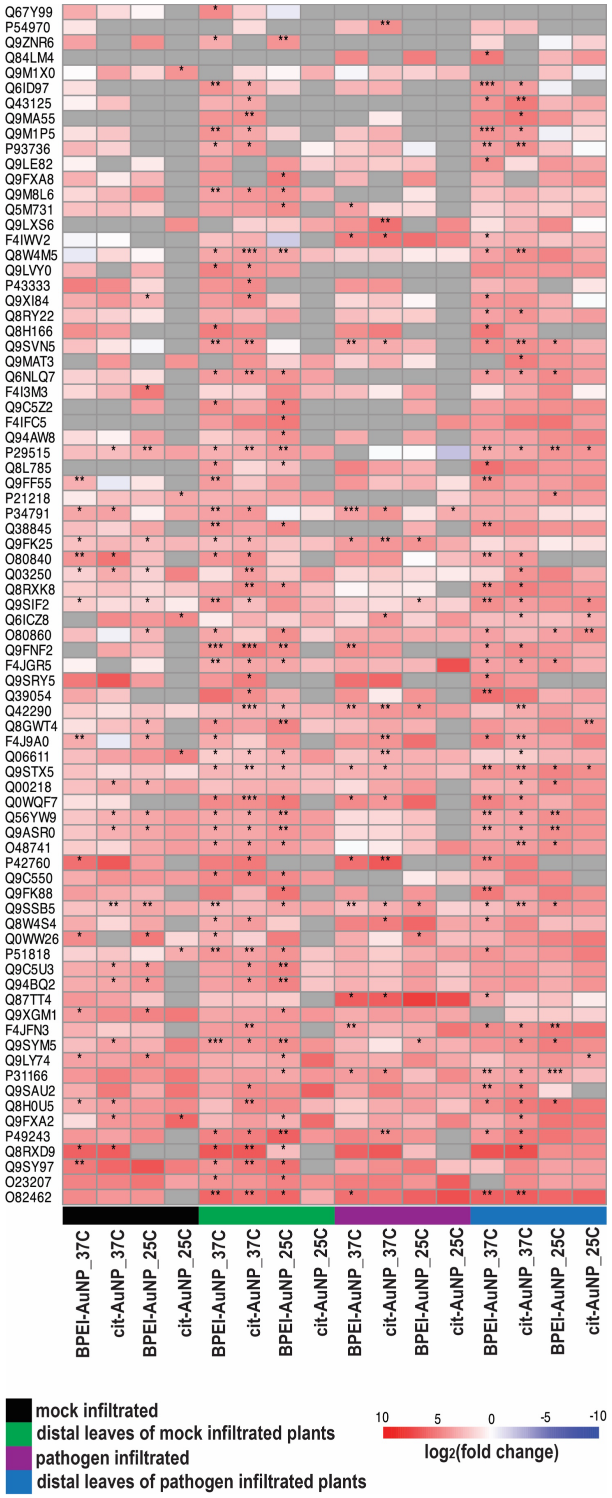

**Supplementary Fig. 17.** Heatmap with hierarchical clustering of a subset of $log_2$(fold change) values, calculated relative to mock samples, for temperature dependent corona proteins. The coronas were formed in mock infiltrated, pathogen infiltrated, and respective distal leaf lysates at 37°C and 25°C. Proteins were filtered to plot $log_2(fc) > 4$ and $p$-values $< 0.05$ observed in at least one sample. Grey boxes signify incalculable fold change. Asterisks denote significance levels (*, $p < 0.05$; **, $p < 0.01$; ***, $p < 0.001$).

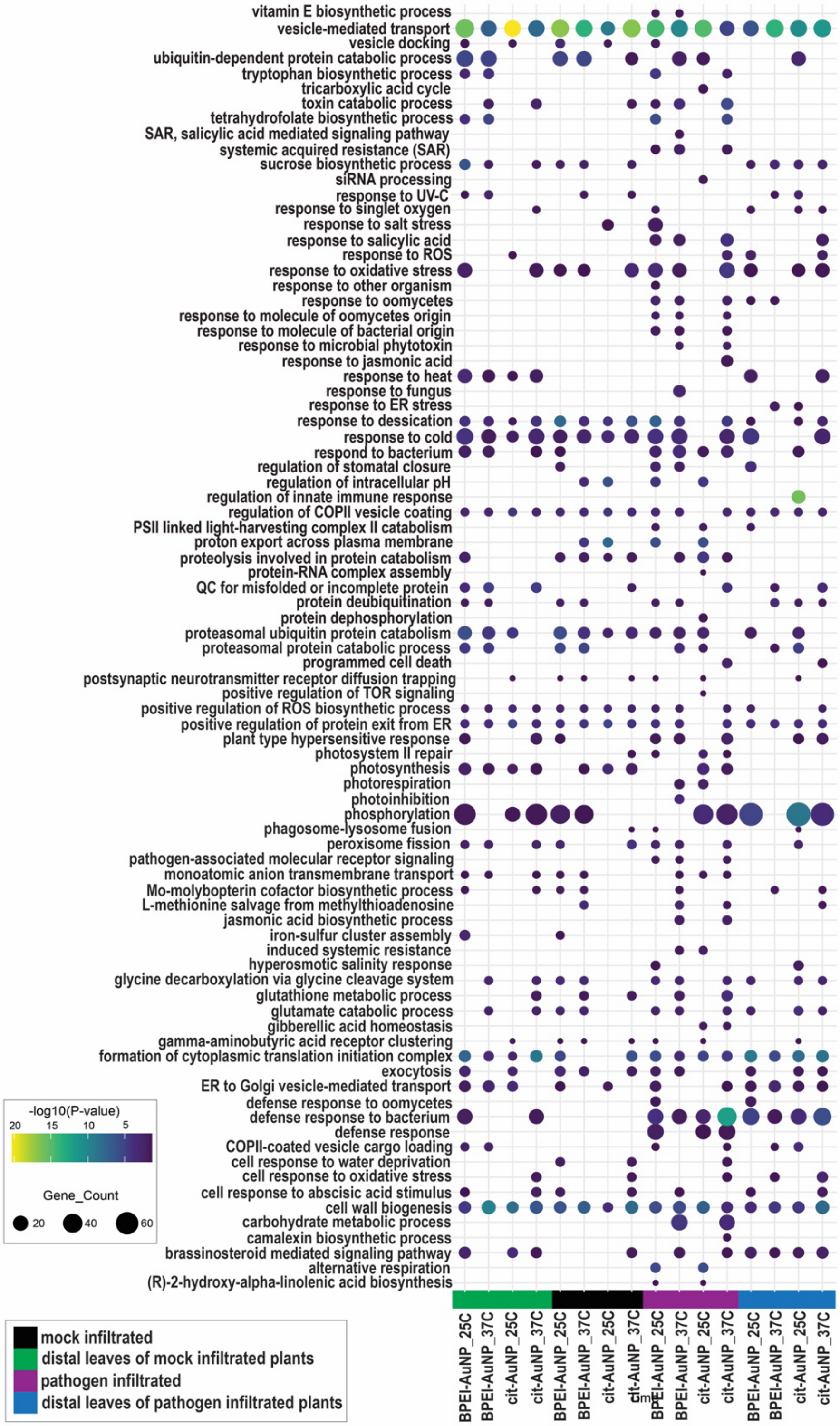


**Supplementary Fig. 18.** Temperature dependence on enriched biological processes associated with 'unique' corona proteins. The x-axis consists of the temperature-dependent samples analyzed by nano-omics. Names of the enriched biological processes are shown on the y-axis. Gene counts are illustrated by dot size, significance is depicted with a color scale of -log(*p*-values), and sample types are color coded.

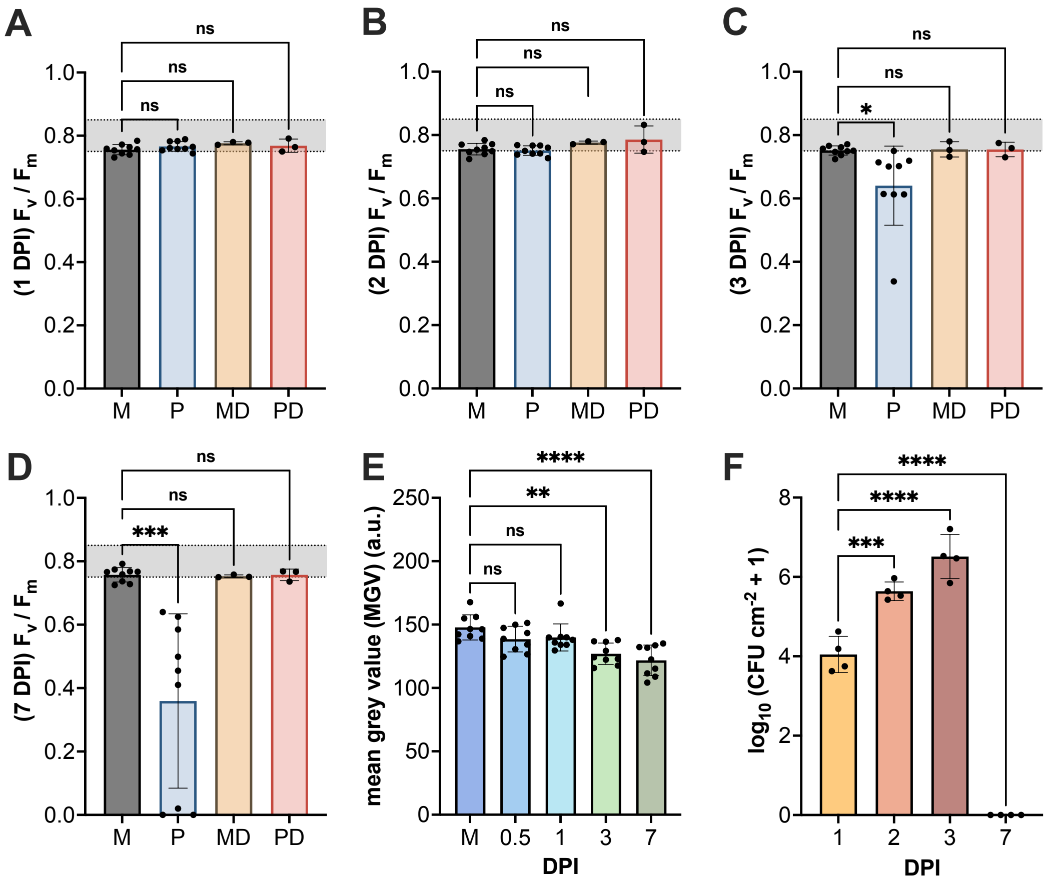


**Supplementary Fig. 19.** Conventional assessments of plant stress. Photosynthetic activity was analyzed by measuring chlorophyll fluorescence of photosystem II in a dark adapted state ($F_v/F_m$) after **A)** 1-, **B)** 2-, **C)** 3-, and **D)** 7-DPI with *P. syringae* or mock solution. Photosynthetic activity was measured using a MINI-PAM-II fluorometer, which measures light excited chlorophyll fluorescence.[1,2] Both directly infiltrated leaves (*n*=9) and distal leaves (*n*=3) were measured with the chlorophyll fluorometer. M: mock; P: pathogen infiltrated; MD: mock distal leaf; PD: distal leaf of pathogen infiltrated plant. The shaded region between 0.75–0.85 represents the values for "unstressed" healthy *A. thaliana*.[3] **E)** Densiometric analysis of plant photographs (Supplementary Fig. 2) was used to quantify the extent of stress and/or infection. Three biological replicates were used for the mock and each timepoint; regions of interest (ROI) were drawn around 3 leaves from each replicate (*n*=9) and analyzed by measuring the mean grey value (MGV) of the ROI. M: mock. **F)** Quantification of the bacterial recovery from infected *A. thaliana* at multiple time points (*n*=4). The mean ± s.d. is shown in all plots and statistical analysis was conducted with an ordinary one-way ANOVA and Tukey's multiple comparisons test.

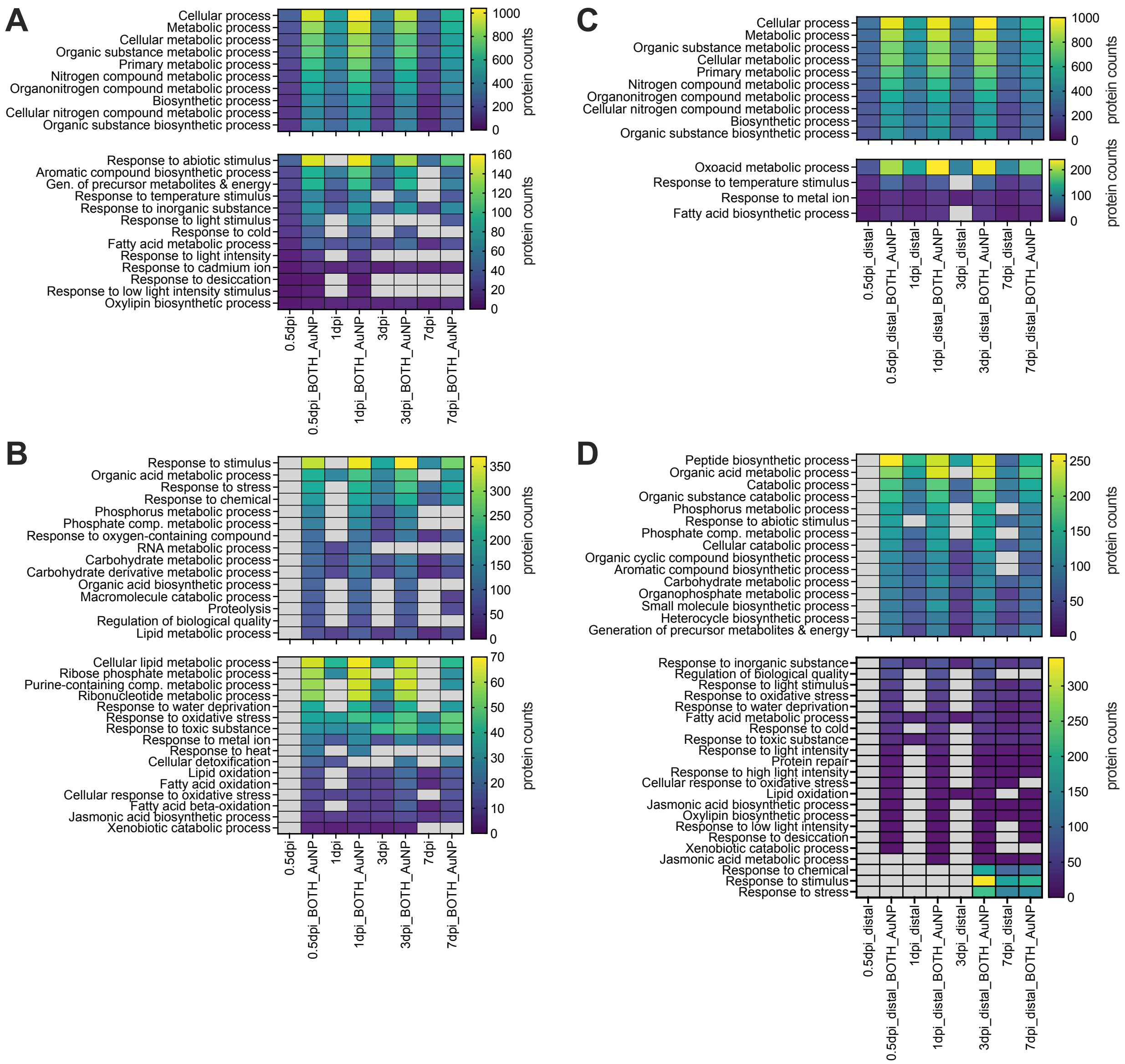


**Supplementary Fig. 20.** Gene ontology analysis of enriched biological processes associated with upregulated/enriched and unique proteins from lysates and AuNP coronas conducted with the STRING database.[4] The heatmaps show the quantity of proteins involved in processes that **A)** include and **B)** exclude processes enriched in 0.5-DPI lysates, and **C)** include and **D)** exclude processes enriched in 0.5-DPI distal lysates. All heatmap legends indicate the range of protein counts associated with each biological process.

| | Conventional approach | | | | | | | | | | |
|---|---|---|---|---|---|---|---|---|---|---|---|
| | Control | mock | mock distal | 0.5 DPI | 1 DPI | 3 DPI | 7 DPI | 0.5 DPI distal | 1 DPI distal | 3 DPI distal | 7 DPI distal |
| avg | 1278.3 | 908.7 | 1131.0 | 915.0 | 1282.3 | 1203.3 | 937.0 | 877.7 | 1254.0 | 956.7 | 1126.3 |
| std | 89.0 | 163.2 | 36.4 | 81.5 | 82.2 | 192.6 | 333.5 | 254.6 | 67.2 | 120.0 | 96.0 |
| cv | 7.0% | 18.0% | 3.2% | 8.9% | 6.4% | 16.0% | 35.6% | 29.0% | 5.4% | 12.5% | 8.5% |

| | Nano-omic approach (BPEI-AuNP) | | | | | | | | | | |
|---|---|---|---|---|---|---|---|---|---|---|---|
| | Control | mock | mock distal | 0.5 DPI | 1 DPI | 3 DPI | 7 DPI | 0.5 DPI distal | 1 DPI distal | 3 DPI distal | 7 DPI distal |
| avg | 1504.0 | 1213.0 | 1031.7 | 1367.0 | 1533.0 | 1574.7 | 1292.0 | 1362.3 | 1355.3 | 1535.3 | 1146.7 |
| std | 83.1 | 112.6 | 97.7 | 126.0 | 49.5 | 224.8 | 164.1 | 188.4 | 48.8 | 63.1 | 94.9 |
| cv | 5.5% | 9.3% | 9.5% | 9.2% | 3.2% | 14.3% | 12.7% | 13.8% | 3.6% | 4.1% | 8.3% |

| | Nano-omic approach (cit-AuNP) | | | | | | | | | | |
|---|---|---|---|---|---|---|---|---|---|---|---|
| | Control | mock | mock distal | 0.5 DPI | 1 DPI | 3 DPI | 7 DPI | 0.5 DPI distal | 1 DPI distal | 3 DPI distal | 7 DPI distal |
| avg | 957.7 | 855.3 | 1143.7 | 1418.7 | 1429.3 | 1421.7 | 1227.3 | 1286.7 | 1285.7 | 1297.3 | 475.0 |
| std | 360.0 | 229.6 | 111.1 | 120.2 | 111.7 | 285.2 | 89.8 | 159.5 | 37.4 | 155.0 | 89.2 |
| cv | 37.6% | 26.8% | 9.7% | 8.5% | 7.8% | 20.1% | 7.3% | 12.4% | 2.9% | 11.9% | 18.8% |

**Supplementary Table 1.** The averages (avg) of identified proteins with their respective standard deviations (std) and CV percentages (cv) for each sample type ($n$ = 3).
